# Modeling clonal evolution and oncogenic dependency in vivo in the context of hematopoietic transformation

**DOI:** 10.1101/2022.05.18.492524

**Authors:** Robert L. Bowman, Andrew Dunbar, Tanmay Mishra, Wenbin Xiao, Michael R. Waarts, Inés Fernández Maestre, Shira E. Eisman, Louise Cai, Sheng F. Cai, Pablo Sanchez Vela, Shoron Mowla, Anthony R. Martinez Benitez, Young Park, Isabelle S. Csete, Aishwarya Krishnan, Darren Lee, Nayla Boorady, Chad R. Potts, Matthew T. Jenkins, Martin P. Carroll, Sara E. Meyer, Linde A. Miles, P. Brent Ferrell, Jennifer J. Trowbridge, Ross L. Levine

**Author notes:** Corresponding Author: Dr. Ross Levine.

## Abstract

Cancer evolution is a multifaceted process involving the acquisition of somatic mutations and progressive epigenetic dysregulation of cellular fate. Both cell-intrinsic mechanisms and environmental interactions provide selective pressures capable of promoting clonal evolution and expansion, with single-cell and bulk DNA sequencing offering increased resolution into this process^1-4^. Advances in genome editing, single-cell biology and expressed lentiviral barcoding have enabled new insights into how transcriptional/epigenetic states change with clonal evolution^5,6^. Despite the extensive catalog of genomic alterations revealed by resequencing studies^7,8^, there remain limited means to functionally model and perturb this evolutionary process in experimental systems^9^. Here we integrated multi-recombinase (Cre, Flp, and Dre) tools for modeling reversible, sequential mutagenesis from premalignant clonal hematopoiesis to acute myeloid leukemia. We demonstrate that somatic acquisition of *Flt3* activating mutations elicits distinct phases of acute and chronic activation resulting in differential cooperativity with *Npm1* and *Dnmt3a* disease alleles. We next developed a generalizable allelic framework allowing for the reversible expression of oncogenic mutations at their endogenous loci. We found that reversal of mutant *Flt3* resulted in rapid leukemic regression with distinct alterations in cellular compartments depending upon co-occurring mutations. These studies provide a path to model sequential mutagenesis and deterministically investigate mechanisms of transformation and oncogenic dependency in the context of clonal evolution.

## MAIN

Gene discovery and genomic landscape studies have elucidated a spectrum of somatic mutations in human cancers, with progressive changes in clonal fitness through iterative cycles of selection, expansion, and stochastic drift^4,10^. Large scale studies on clinical isolates have observed intratumoral heterogeneity through spatially/temporally separated sampling^11-13^ or through targeted single-cell DNA-sequencing studies^2,3^. In myeloid malignancies, mutations in epigenetic modifiers (e.g. *DNMT3A, TET2*) have been identified as subclonal, low variant allele frequency (VAF) events in the blood of healthy individuals in premalignant clonal hematopoiesis (CH) and as clonal events in leukemia. Despite a paucity of *FLT3* mutations in CH, the receptor tyrosine kinase *FLT3* is the most commonly mutated gene in acute myeloid leukemia (AML) often presenting as a relatively late hit in leukemic evolution. Patients harboring the triplet of *DNMT3A, NPM1* and *FLT3* mutations suffer a dismal prognosis^7^ and represent a genetic archetype for the transition from premalignant CH to transformed AML. Our understanding of the processes underlying this evolution is limited by a lack of deterministic experimental systems to evaluate the consequences of sequential mutational acquisition *in vivo*.

Experimental systems have largely relied upon ectopic overexpression models, Cre-Lox technology^14^, and the increasing use of CRISPR/CAS9 genome editing in different species^15-18^. Questions regarding sequential mutagenesis and contextual cooperativity are not adequately addressed by these existing systems, particularly for mutant *FLT3 ^19^* where internal tandem duplication (ITD) mutations occur late in leukemic transformation yet have not been investigated when acquired subsequent to other leukemia disease alleles. Here we model stepwise clonal evolution in myeloid malignancy by integrating new and existing tools centered on the orthogonal DNA recombinases Cre, Dre, and Flp (**Extended Data Fig. 1a**). We developed inducible models of *Flt3*^ITD^ for both Flp and Dre recombinase which, when paired with cooperating alleles, result in lethal, penetrant models of AML. We further deploy methods for orthogonally inducing Cre and Dre recombinase enabling extended sequential mutagenesis and for investigating the consequences of perturbing cancer evolution through reversible oncogene expression.

### Inducible mouse model of Flt3 internal tandem duplication

We first developed an endogenously targeted, Flp-inducible *Flt3*^Frt-ITD^ allele which could be inverted by a tamoxifen (TAM) inducible FlpoERT2 allele (**Figure 1a, Extended Data Fig. 1b**). By 6 weeks post tamoxifen administration, we observed penetrant leukocytosis (mean WBC: *Flt3*^Frt-ITD^ 38.21, *WT* 10.37, p ≤ 4.5×10^-4^) driven by a myeloid bias and expansion of cKit+ cells in the blood (mean 6w:4.8% vs. Control: 0.31% p ≤ 5.27×10^-3^**; Figure 1b, Extended Data Fig. 1b-c).** By 8-10 weeks, this leukocytosis had largely resolved, with WBCs decreasing to near normal levels (*p* ≤ 0.133) despite persistent anemia (mean HCT%: 40.4 vs. 51.4, *p* ≤ 1.71×10^-7^), and thrombocytopenia (mean PLT K/μL: 467 vs. 1240, *p* ≤ 8.96×10^-6^**; Extended Data Fig. 1d**). *Flt3*^Frt-ITD^ mice developed significant splenomegaly (mean: 361.99mg vs. 90.35mg *p* ≤ 2.84×10^-3^), and a pathological myeloid infiltrate into both the spleen and liver (**Figure 1c**, **Extended Data 1e)**. An increase in immature myeloid cells, reduction in megakaryocytes, and near absence of erythroid cells was evident in the bone marrow in *Flt3^Frt-ITD^* mice. Splenic architecture was also disrupted— pathological findings which persisted even after resolution of leukocytosis (**Figure 1c, Extended Data Fig. 1f)**. Along this time course, we observed an expansion of lineage negative hematopoietic stem and progenitor cells (HSPCs; mean 6w 50.1% vs. Control 8.55%, *p* ≤ 0.0016) specifically the Lin^-^Sca-1^+^cKit^+^ (LSKs) in the bone marrow (4w vs. Control marrow: *p ≤* 0.0382; **Figure 1d, Extended Data Fig. 1g)**.

**Figure 1:**
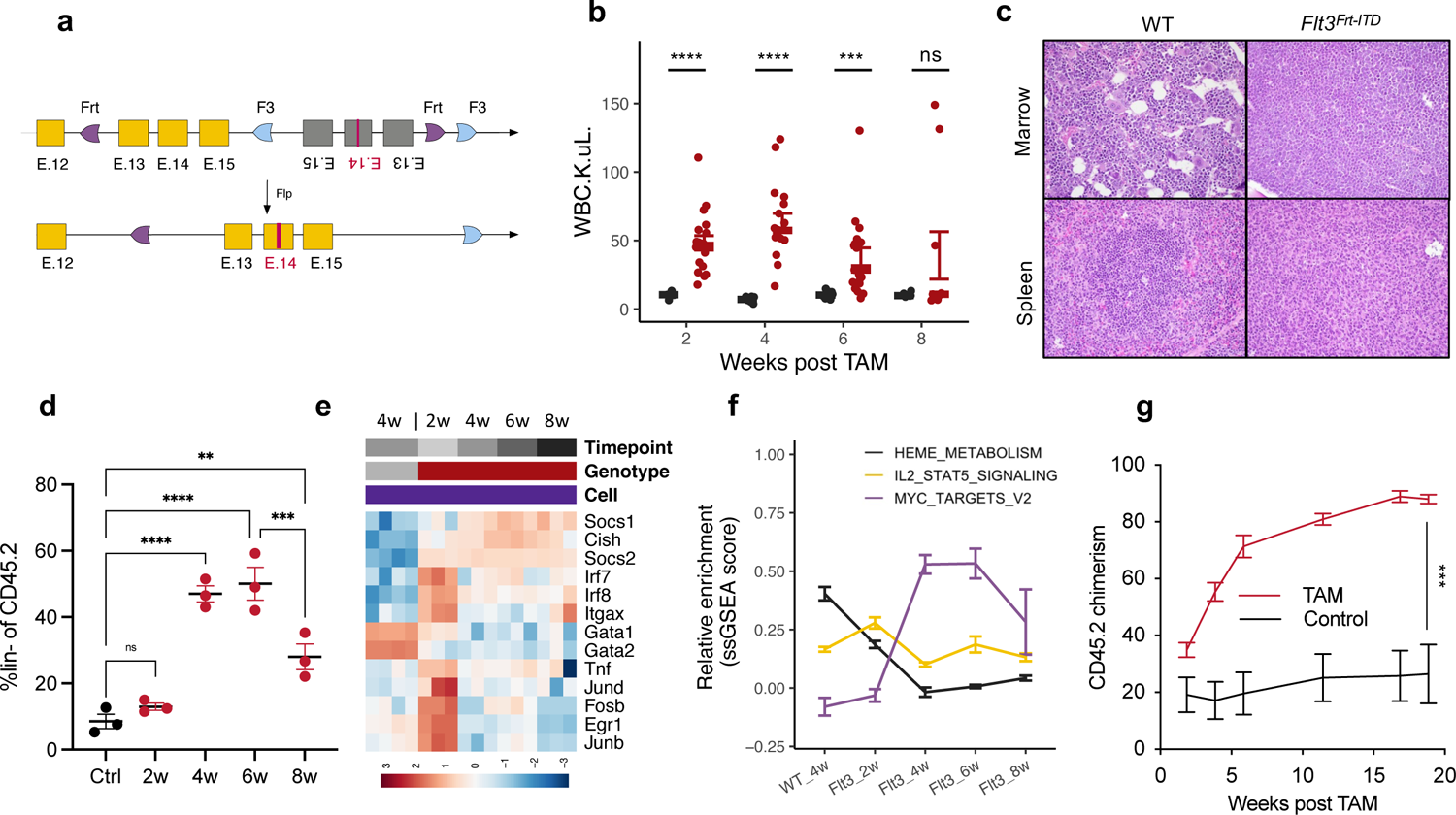
Flp-recombinase inducible Flt3^ITD^ activation. **a,** schematic *Flt3^Frt-ITD^* targeted to the endogenous *Flt3* locus replacing WT exons. Flp-recombination deletes WT exons and inverts mutant exons resulting in *Flt3*^ITD^ expression. **b,** strip chart of WBC K/μL in WT (black) or Rosa26:FlpoERT2 *Flt3^Frt-ITD^* (red) mice at the indicated timepoint posts TAM. (n=10-19 per group) **c,** H&E staining of bone marrow or spleen in WT or Rosa26:FlpoERT2 *Flt3^Frt-ITD^* mice 16 weeks post TAM treatment (400x). **d,** strip chart indicating %lin^-^ cells in WT (Ctrl) or *Flt3^Frt-ITD^* bone marrow. (red; n=3 per group). **e,** Row normalized heatmap of RNA-sequencing data in WT (grey) or *Flt3^Frt-ITD^* (red) LSKs following TAM treatment at the indicated timepoints. **f,** Z-scored ssGSEA values for the indicated Hallmark geneset in either WT or *Flt3^Frt-ITD^* mice following TAM treatment. (n=3 per group). **g,** Peripheral blood chimerism (%Cd45.2) in mice engrafted with WT (Cd45.1) and *Flt3^Frt-ITD^* (Cd45.2) cells following treatment with TAM (red) or control (black). (n=6-7 per group). Error bars reflect the mean ± s.e.m.; and p-values are calculated by Student’s t-test (b,g) and Fisher’s LSD test (d). *\*\* p ≤ 0.01* *** *p ≤ 0.001* **** *p ≤ 0.0001*

We next wanted to determine how somatic acquisition of *Flt3*^Frt-ITD^ mutations changed the transcriptional landscape of HSPCs. We performed low pass, multiplexed 3’ RNA-sequencing on purified LSKs and granulocytic-monocytic progenitors (GMPs; cKit^+^Sca1^-^Cd34^+^Cd16/32^+^) from *Flt3^ITD^* mice 2, 4, 6 or 8 weeks after TAM administration and from Flt3^WT^ mice 4 weeks after TAM **(Extended Data Fig. 2a)**. Along this time course >200 genes were downregulated in LSKs with *Flt3*^Frt-ITD^ activation, and >500 were upregulated with at least one time point **(Extended Data Fig. 2b, Supplementary Table 1a)**. These alterations could be largely consolidated into three distinct phases of transcriptional alterations along the 8-week time course including genes which were (1) acutely (in)activated and dysregulation was sustained (2) delayed changes in gene expression seen after 4+ weeks, and (3) transiently dysregulated at 2 weeks post *Flt3^ITD^* activation and then normalized **(Extended Data Fig. 2c)**. Sustained changes in transcriptional output included reduced expression of heme synthesis genes (*FDR ≤* 2.05×10^-6^) with concomitant downregulated expression of the erythroid transcription factors *Gata1* (8w *p* ≤ 1.51×10^-16^) and *Gata2* (8w *p* ≤ 7.11×10^-23^; **Figure 1e-f)** and increased expression of MYC target genes (*FDR ≤* 4.97×10^-3^). Expression of STAT5 target genes peaked two weeks following mutational activation (FDR ≤ 0.044) whereas increased expression of negative signaling regulators including *Cish* (8w vs. WT *p* ≤ 8.99×10^-15^; log2FC 2.82) and *Socs2* persisted (8w vs WT *p* ≤ 8.17e-133; log2FC 4.1; **Figure 1e-f; Extended Data Fig. 2d)**. In addition to transient increased expression of dendritic-cell like genes at 2 weeks, gene expression signatures indicative of MAPK/KRAS signaling were also upregulated (*FDR* ≤ 0.00389) with transient expression of AP-1 complex members (*Jund, Junb, Fos*) and the MAPK target *Egr1* (**Figure 1e-f; Extended Data Fig. 2e)**. Critical regulators of hematopoietic stem cell self-renewal including *Tal1, Meis1, Mecom, Hoxb5,* and *Hoxa9* were consistently decreased in expression **(Extended Data 2f)**.

To understand the functional consequences of somatic *Flt3*^Frt-ITD^ activation on stem/progenitor function and relative fitness, we performed competitive transplantation between *Flt3*^Frt-ITD^ and Cd45.1 WT unfractionated marrow. Activation of *Flt3*^Frt-ITD^ with TAM 4 weeks after transplantation resulted in robust, sustained competitive outgrowth of *Flt3*^Frt-ITD^ cells compared to WT competitor cells (mean Cd45.2% at 18w 88.89% vs. 25.8%, *p* ≤ 6.54×10^-4^; **Figure 1g)**. By contrast, *Flt3*^Frt-ITD^ mutant cells were incapable of robustly engrafting into recipient mice and propagating disease in secondary recipients **(Extended Data Fig. 2g)**. Consistent with the lack of self-renewal *in vivo*, *Flt3*^Frt-ITD^ acquisition resulted in the depletion of long-term hematopoietic stem cells (LT-HSCs; Lin^-^Sca-1^+^cKit^+^CD150^+^CD48^-^), both with respect to Cd45.2 *Flt3*^Frt-ITD^ (mean±s.e.m.; Control: 1.37±0.22% vs. TAM: 0.024±0.007%, *p* ≤ 0.0004) and cell non-autonomous depletion of Cd45.1 WT stem cells (mean±s.e.m.; Control: 0.9±0.2% vs. TAM: 0.18±0.1, *p* ≤ 0.0194; **Extended Data Fig. 2h-i)**, resolving a question unanswered by existing tools ^20,21^. These findings demonstrate that while *Flt3*^Frt-ITD^ cells can give rise to a robust myeloproliferative disease, HSPCs lose long-term hematopoietic self-renewal once *Flt3*^ITD^ is acquired which we hypothesize represents a key featured conferred by cooperating AML disease alleles.

### Cooperating mutations license mutant-*Flt3* to promote leukemic transformation

We next crossed *Flt3*^Frt-ITD^ mice with a *Npm^Frt-c^* mutant allele^9^ and somatically activated both alleles with a TAM-inducible FlpoERT2 allele. In contrast to primary *Flt3*^Frt-ITD^ mice which had a median survival of 48 weeks post-TAM, *Npm^Frt-c^*-*Flt3*^Frt-ITD^ (NF) mice rapidly succumbed to leukemia within 4 weeks following TAM administration (log rank *p* ≤ 8.78×10^-7^; **Figure 2a; Extended Data Fig. 3a).** In a transplantation model, mice engrafted with *Npm^Frt-c^*-*Flt3*^Frt-ITD^ cells exhibited a robust increase in WBC (mean 380.6 K/μL, ANOVA *p*≤ 0.0001) with rapidly progressive anemia (mean HCT 15.99% ANOVA, *p*≤ 0.0001) and thrombocytopenia (mean PLT 97.40K/μL, ANOVA *p*≤ 0.0001) by 4 weeks post-transplantation which was not seen with somatic expression of either *Flt3*^Frt-ITD^ or *Npm^Frt-c^* alone. All recipient mice showed pathological features consistent with a fully penetrant, rapid AML (**Figure 2b; Extended Data Fig. 3b-c)**. Unlike *Flt3*^Frt-ITD^ single mutant mice, the increase in WBC with *Npm^Frt-c^*-*Flt3*^Frt-ITD^ did not normalize and continued to increase to levels observed in AML patients with *FLT3/NPM1^c^* mutations,^22^ even with limiting doses of TAM exposure, which reduced the proportion of mutant cells which were induced **(Extended Data Fig. 3d)**.

**Figure 2:**
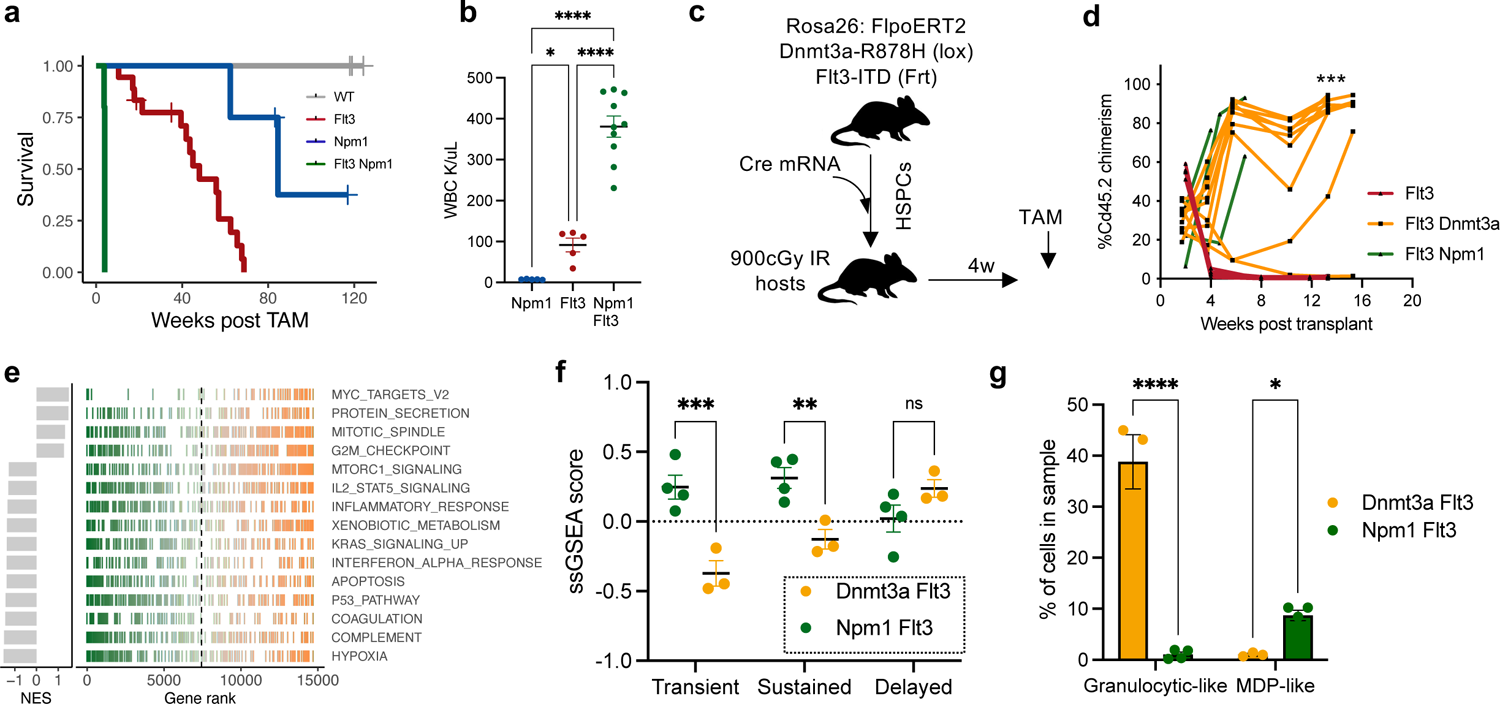
Flt3^ITD^-driven models of leukemogenesis. **a,** Kaplan-Meier survival curve for Rosa26:FlpoERT2 WT (grey), *Flt3^Frt-ITD^* (red), *Npm1^c-Frt^* (blue), or *Npm1^c-Frt^ Flt3^Frt-ITD^* (green) following treatment with TAM. (n=4-18 per group). **b,** Bone marrow was transplanted into lethally irradiated recipient mice. After transplant (4w) mice were treated with TAM to activate the indicated genotypes. Strip chart indicates WBC. (n=5-10 per group) **c,** Bone marrow transplant-mediated sequential mutagenesis experimental schematic indicating order of *Dnmt3a^Loox-R878H^* (Cre mRNA) and *Flt3^Frt-ITD^* (TAM) activation. **d,** Bone marrow from Cd45.1 WT or Cd45.2 cells from *Npm1^c-Frt^ Flt3^Frt-ITD^* (green), *Dnmt3a^Loox-R878H^ Flt3^Frt-ITD^* (orange) *or Flt3^Frt-ITD^* only (red) mutant mice >20w after TAM treatment were transplanted in competition into secondary lethally irradiated recipients. Peripheral blood chimerism (%Cd45.2) depicted for individual mice. (n=3-10 per group). **e,** RNA-sequencing data from *Npm1^Lox-C^ Flt3^GL-ITD^* or *Dnmt3a^Loox-R878H^ Flt3^Frt-ITD^* control LSKs. Gene ranks from GSEA are depicted as a linechart for the indicated HALLMARK gene set. **f,** normalized ssGSEA scores on RNA-sequencing data from LSKs sorted from symptomatic *Npm1^c-Frt^ Flt3^Frt-ITD^* (green) or *Dnmt3a^Loox-R878H^ Flt3^Frt-ITD^* (orange) mice. Genesets were derived from Extended Data 2c. (n=3 per group) **g,** Barplot depicting %bone marrow cells with a Granulocyte-like (left, Cluster 1) or MDP-like (right, Cluster 11) immunophentype characterized by CyTOF. (n=3-4 per group). Error bars reflect the mean ± s.e.m.; and p-values are calculated by Fisher’s LSD test (b,f,g) and Student’s t-test (d) and *\* p ≤ 0.05 ** p ≤ 0.01* *** *p ≤ 0.001* **** *p ≤ 0.0001*

We next sought to assess the impact of mutant *Flt3* acquisition subsequent to a common co-occurring CH mutation, specifically *Dnmt3a^Lox-R878H^*. We administered Cre mRNA by electroporation *ex vivo* to induce *Dnmt3a^Lox-R878H^* expression, and TAM was used to activate *Flt3*^Frt-ITD^ four weeks after transplant (**Figure 2c)**. While the *Npm1^Frt-c^*-*Flt3*^Frt-ITD^ mice rapidly succumbed to AML with *Flt3*^Frt-ITD^ activation, there were no significant differences in survival between *Dnmt3a^Lox-R878H^*-*Flt3*^Frt-ITD^ and *Flt3*^Frt-ITD^ only mice (median survival DF 25.5w vs. F 25w, log rank *p* ≤ 0.69; **Extended Data Fig. 3e)**. In these *Dnmt3a^Lox-R878H^-Flt3*^Frt-ITD^ (DF) mice, we observed the same increase and subsequent decreases in WBC as in the *Flt3*^Frt-ITD^ alone model (mean DF WBC: 8w 41 K/μL vs. 13w 20 K/μL, *p* ≤ 0.0015); however, by 18-20 weeks post-TAM, the DF mice developed progressive leukocytosis (mean WBC 110 K/μL, *p* ≤ 0.0269**; Extended Data Fig. 3f)**. Critically, like *Npm1^Frt-c^*-*Flt3*^Frt-ITD^, *Dnmt3a^Lox-R878H^*-*Flt3*^Frt-ITD^ cells were capable of engrafting into secondary recipients (mean Cd45.2% 12w DF 76.3% vs. F 0.99%, *p* ≤ 1.17×10^-4^) and propagating disease, albeit at a longer latency than the aggressive *Npm1^Frt-c^*-*Flt3*^Frt-ITD^ model (**Figure 2d)**.

Gene expression analysis revealed that in comparison to *Npm1^Frt-c^*-*Flt3*^Frt-ITD^ LSKs, *Dnmt3a^Lox-R878H^*-*Flt3*^Frt-ITD^ LSKs were enriched for expression of E2F targets (GSEA NES 1.74, *FDR* ≤ 8.43×10^-5^), cell cycle signatures (GSEA FDR 1.46 ≤ 2.96×10^-3^) and expression of immature markers including *Gpr56*, *Cd34*, *Abcc1*, *Kit* and *Mpl* (**Figure 2e; Supplementary Table 1b)**. In contrast, *Npm1^Frt-c^*-*Flt3*^Frt-ITD^ leukemias expressed lower levels of Cd34 within the stem cell compartment (*p* ≤ 0.0032), matching the clinical immunophenotype ^23,24^, and had higher levels of *Hox* gene expression particularly *Hoxa7* and *Hoxa9* **(Extended Data Fig. 3g-h; Supplementary Table 1c)**. LSKs and GMPs from *Npm1^Frt-c^*-*Flt3*^Frt-ITD^ leukemias had increased inflammatory gene expression particularly for TNFα and NFκB, reminiscent of early-stage *Flt3*^Frt-ITD^ activation. In particular, *Npm1^Frt-c^*-*Flt3*^Frt-ITD^ LSKs were specifically enriched for genes transiently activated with *Flt3*^Frt-ITD^ acquisition alone (LSK *p* ≤ 0.0001) suggesting that *Npm1^c^* cooperates to “lock” *Flt3^ITD^* cells into an early activation state (**Figure 2f)**.

Immunophenotypically, we observed an expansion of the progenitor compartment for *Dnmt3a^Lox-R878H^*-*Flt3*^Frt-ITD^ cells (%GMP of Cd45.2 mean±s.e.m. DF: 7.1±1.16% vs. NF: 2.32±0.41%, *p* ≤ 0.0029) while *Npm1^Frt-c^*-*Flt3*^Frt-ITD^ leukemias were enriched for LSKs (%LSK of Cd45.2 mean±s.e.m. DF: 0.4 ± 0.09 vs NF: 1.8±0.16, *p* ≤ 0.0001**; Extended Data Fig. 4a)**. Mass cytometry (CyTOF) analysis showed that *Npm1^Frt-c^*-*Flt3*^Frt-ITD^ leukemias had increased numbers of cells enriched for dendritic-like characteristics (expression of Cd11c, *p* ≤ 1.46×10^-3^), while *Dnmt3a^Lox-R878H^*-*Flt3*^Frt-ITD^ leukemias had expression of markers more consistent with a granulocytic bias (*p* ≤ 3.84×10^-4^; **Figure 2g; Extended Data Fig. 4b-d)**. These differentiation biases were present even at the level of stem cells, with LSKs from *Npm1^Frt-c^*-*Flt3*^Frt-ITD^ mice expressing higher levels of the GM-CSF receptor *Csf2ra* (*p* ≤ 2.08×10^-3^) and *Dnmt3a^Lox-R878H^*-*Flt3*^Frt-ITD^ LSKs expressing increased levels of the G-CSF receptor *Csf3r* (*p* ≤ 7.39×10^-8^) **(Extended Data Fig. 3h)**. Collectively, these models demonstrate that different co-occurring mutations are capable of transforming *Flt3* mutant cells, albeit with distinct latencies, phases of cooperativity, and transcriptional /immunophenotypic outputs.

### Expanding the sequential mutagenesis toolkit with Dre recombinase

Given most inducible murine models use Cre or Flp recombinase and the vast majority of human cancers have 3 or more pathogenic disease alleles, we sought to develop AML models with 3 orthogonal recombinases, aiming to add reversible mutant activation to our approach. We therefore chose to develop a model induced by Dre recombinase, a Cre-homologue capable of recombining Rox sites. We developed a dual-recombinase strategy to activate and then inactivate a mutation of interest termed GOLDI-Lox (<u>g</u>overning <u>o</u>ncogenic loci by <u>d</u>re inversion and <u>lox</u> deletion; **Figure 3a)**. In this model, a pair of Lox2272 sites flank a staggered, heterotypic Rox12 - RoxP pair, which themselves flank an inverted mutant exon encoding the same ITD mutation as in the *Flt3^Frt-ITD^* allele. In the non-recombined state, the proximity (82bp) of the Lox2272 sites prevents Cre recombination at this locus^25^. Upon Dre activation, the *Flt3^ITD^* encoding exons are inverted resulting in reorientation of the internal Lox2272 site and relocation at a sufficient distance from the distal Lox2272 site to permit recombination upon treatment with Cre. A similar strategy was used to reversibly activate *Jak2V617F* in the accompanying manuscript (Dunbar A. & Bowman R. *et. al*, 2022). To evaluate the efficacy of this strategy, we crossed this mouse (*Flt3*^GL-ITD^) to a Dre Reporter (RLTG; TdTomato reporter) and a TAM-inducible Ubc:CreER allele. HSPCs from these mice were infected with a Dre retrovirus for 48 hours with or without Cre induction by 4-hydroxy-tamoxifen (4-OHT). PCR demonstrated that Dre activation was indeed capable of inverting the target locus, while subsequent administration of 4-OHT resulted in deletion of the target locus (**Figure 3b)**.

**Figure 3:**
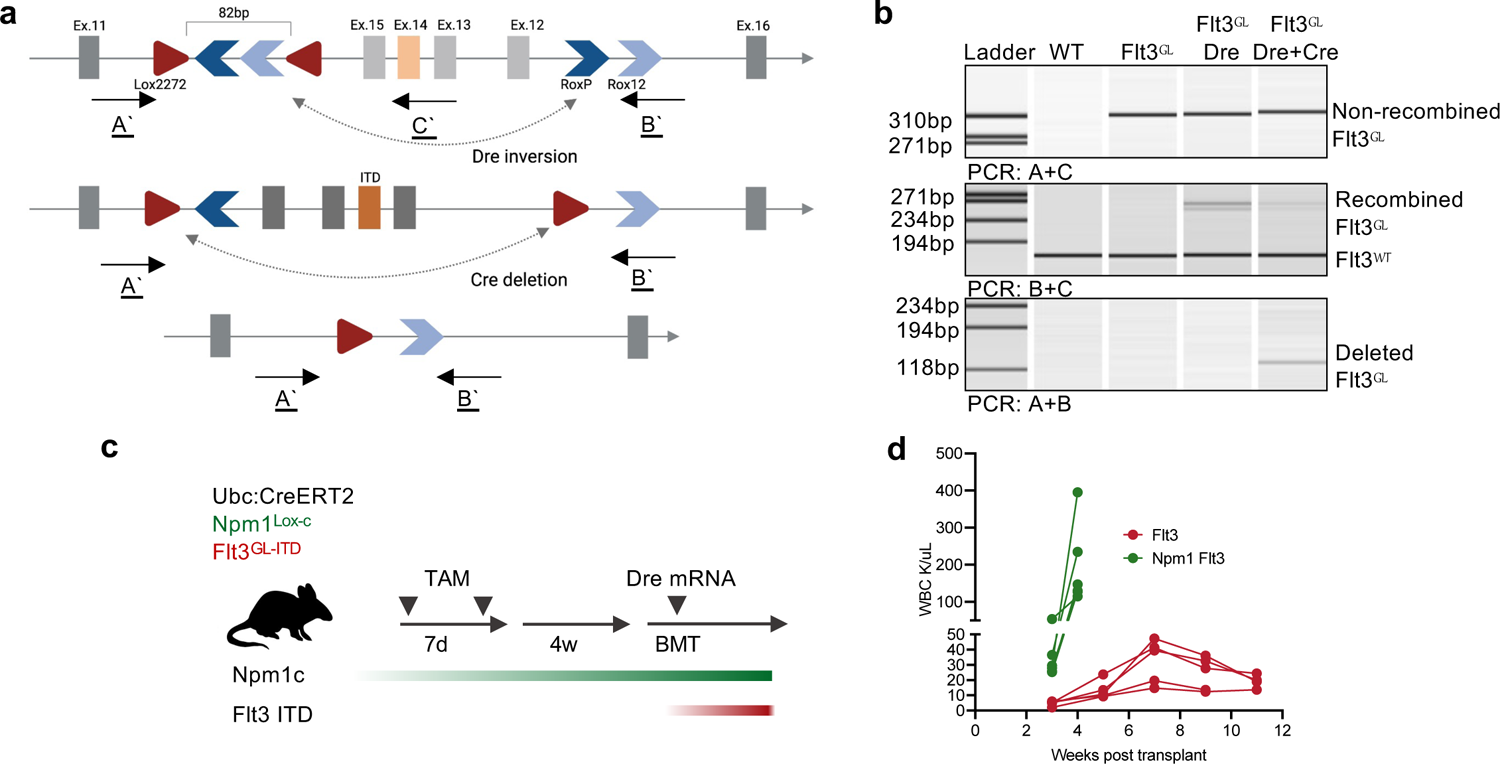
Dual-recombinase, reversible models of oncogene activation. **a,** Schematic depicting GOLDI-Lox knockin *Flt3^GL-ITD^* construct to the endogenous *Flt3* locus, replacing exons 12-15. Flanking exons 11 and 16 are indicated in light grey, Lox2272 sites are indicated by red triangles, heterotypic RoxP and Rox12 sites are indicated in dark and light blue chevrons respectively. Inverted exons 12-15 are present at baseline with the W51 ITD encoded in exon 14 (orange). A’, B’ and C’ indicate relative position of primers used in (b) to detect Dre-mediated inversion and Cre-mediated deletion. **b,** Lin^-^ bone marrow from WT of Ubc:CreERT2 *Flt3^GL-ITD^* mice was infected with MSCV:Dre-IRES-GFP retrovirus or mock infection. Infected cells were treated ± 4-OHT to activate Cre, and DNA was isolated from the whole culture 72 hours post treatment. PCR was used to evaluate Dre-inversion and Cre-deletion, visualized by capillary gel electrophoresis (Qiaxcel). **c,** Schema representing experimental setup where Ubc:CreERT2 *Flt3^GL-ITD^* mice were treated with TAM to activate *Npm1*^Lox-C^. Eight weeks after TAM treatment, mice were euthanized and lin^-^ HSPCs were isolated for Dre mRNA electroporation. Cells (20,000-100,000) were then transplanted into lethally Cd45.1 recipients along with 200,000 whole bone marrow from Cd45.1 mice. Disease development was monitored by complete blood count. **d,** WBC (K/μL) from mice as described in (c). (n=5-6 per group).

To benchmark the *Flt3*^GL-ITD^ allele we crossed it to a Cre-inducible *Npm1^Lox-c^* allele and the TAM-inducible Ubc:CreER. *Npm1^WT^*-*Flt3*^GL-ITD^ or *Npm1^Lox-c^*-*Flt3*^GL-ITD^ mice were treated with TAM, and 8 weeks later, HSPCs were electroporated with Dre mRNA *ex vivo* and then transplanted into lethally irradiated recipient mice (**Figure 3c)**. As expected, mice transplanted with *Npm1^WT^*-*Flt3*^GL-ITD^ mutant cells developed leukocytosis with monocytosis 7w post-transplant (*p* ≤ 0.0103), with a subsequent decrease at 11w post-transplant (*p* ≤ 0.062) (**Figure 3d)**. By contrast, *Npm1^Lox-c^*-*Flt3*^GL-ITD^ cells developed rapid, progressive leukocytosis (4w mean WBC 191.46 K/μL) with fully penetrant progression to AML with similar latency and immunophenotypic characteristics as the Flp-inducible *Npm1^Frt-c^*-*Flt3*^Frt-ITD^ model. Consistent with our previous data in primary human samples showing these mutant alleles potently synergize to promote clonal dominance,^3^ these data underscore that co-occurring mutations with potent mutational synergy can induce AML when activated simultaneously or in sequence.

### Sequential mutagenesis reveals stage and mutant-specific alterations in differentiation

While *Npm1*^C^*-Flt3*^ITD^ combinations potently induce AML in mice, the majority of AML patients with *NPM1*^C^ and *FLT3*^ITD^ mutant alleles acquire these mutations subsequent to an antecedent mutation, most commonly *DNMT3A.* Moreover, the presence of these 3 mutations in concert confers a particularly dismal prognosis for AML patients.^7^ We sought to model this combination of alleles using 3 separably inducible alleles for *Dnmt3a^Lox-R878H^*, *Npm1^Frt-c^*, and *Flt3^GL-ITD^* (**Figure 4a)**. *Dnmt3a^Lox-R878H^* was induced by Cre mRNA electroporation *ex vivo* into HSPCs at transplant. Following transplant, *Npm1^Frt-c^* mutations were then activated in a subset of mice with the TAM-inducible FlpoERT2 allele. By 16 weeks post-*Npm1^Frt-c^* activation, *Dnmt3a^Lox-R878H^*-*Npm1^Frt-c^* (DN) mutant mice developed anemia (M/μL mean±s.e.m. DN 8.2±0.16 vs WT 9.7±0.12, *p* ≤ 0.0001) and a myeloid bias notable for an increase in Cd11b^+^Gr1^high^ granulocytic cells (ANOVA *p* ≤ 0.0014**; Extended Data Fig. 5a)**.

**Figure 4:**
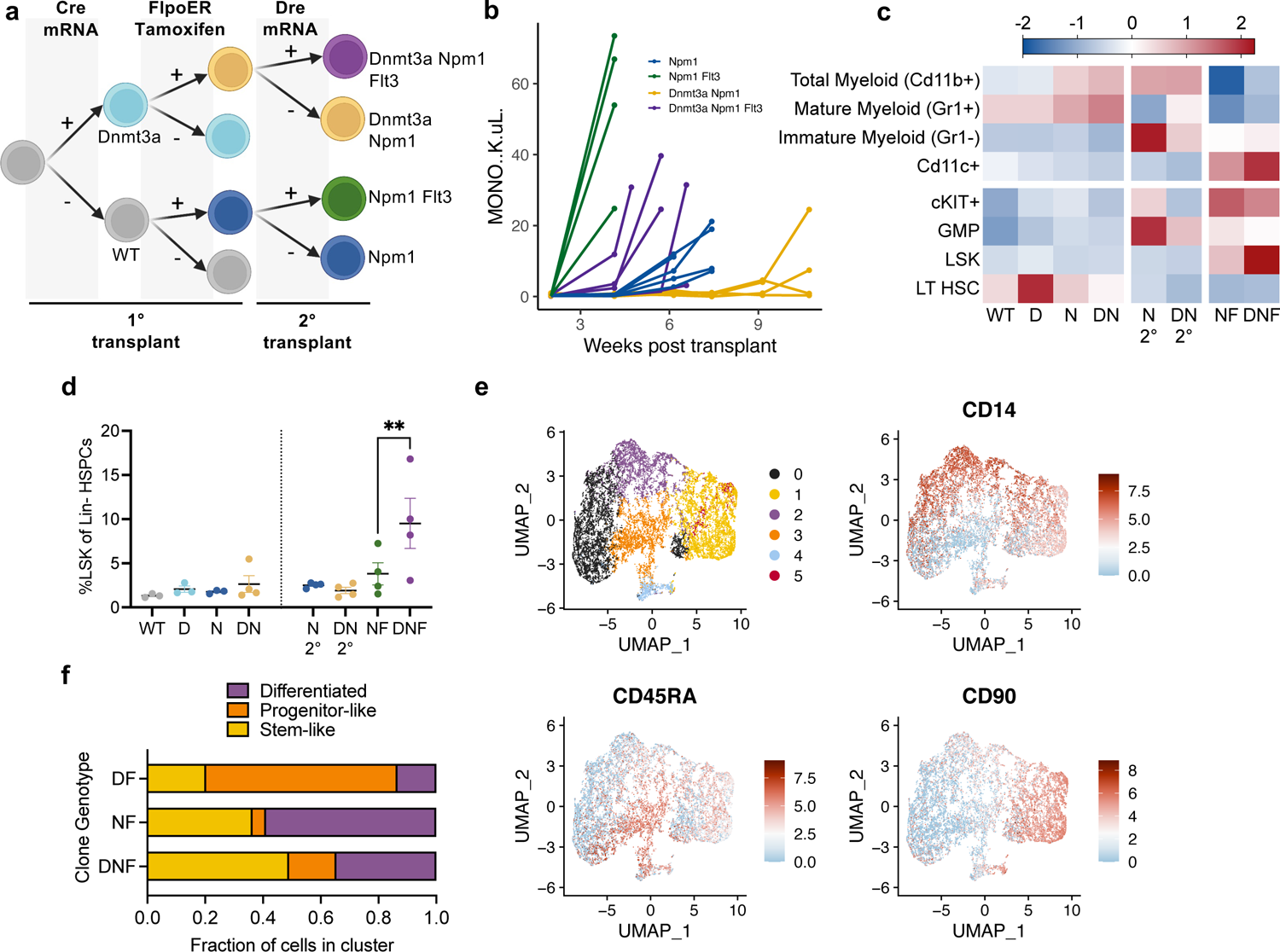
Triple-mutant, sequential models of clonal evolution towards acute myeloid leukemia. **a,** Schematic depicting sequential mutagenesis scheme with *Dnmt3a^Los-R878H^* being activated at first transplant with Cre mRNA, *Npm1^c-Frt^* activated post transplant with TAM-mediated Rosa26:FlpoERT2 activation, and *Flt3^GL-ITD^* finally activated with Dre mRNA at secondary transplant. **b,** Peripheral blood monocyte count (K/μL) in secondary transplant for the indicated genotypes. (n=5 per group). **c,** Heatmap depicting relative abundance of indicated cell populations (rows) for the indicated genotypes (columns) in bone marrow isolated from mice either 20 weeks post transplant (WT, *Dnmt3a^Los-R878H^* (D)*, Npm1^c-Frt^* (N)*, Dnmt3a^Los-R878H^ Npm1^c-Frt^* (DN)) or when symptomatic (*Npm1^c-Frt^* 2° (N 2°)*, Dnmt3a^Los-R878H^ Npm1^c-Frt^* 2° (DN 2°), *Npm1^c-Frt^ Flt3^GL-ITD^* (NF), or *Dnmt3a^Los-R878H^ Npm1^c-Frt^ Flt3^GL-ITD^* (DNF)). Secondary transplant indicated by 2°, following either Dre-mRNA or mock electroporation. **d,** Strip chart from mice depicted in (c) with y-axis indicated %LSKs of Lin^-^ Cd45.2 in bone marrow. (n=3-4 per group). **e,** UMAP of protein abundance on 4 AML patient samples depicting either cluster number (top left), CD14 (top right), CD45RA (bottom left) or CD90 (bottom left) expression. **f,** Stacked barplot indicating fraction of cells present in three of the clusters from (e) for either *DNMT3A-FLT3* (DF), *NPM1-FLT3* (NF) or *DNMT3A-NPM1-FLT3* (DNF) mutant cells. Error bars reflect the mean ± s.e.m.; and p-values are calculated by Fisher’s LSD test (d) *\*\* p ≤ 0.01*.

To model subsequent acquisition of *Flt3^ITD^*, we then electroporated Dre mRNA into an additional subset of *Dnmt3a^Lox-R878H^*-*Npm1^Frt-c^* and *Npm1^Frt-c^* HSPCs followed by transplantation to generate 4 new experimental groups (**Figure 4a)**. *Npm1^Frt-c^*-*Flt3^GL-ITD^* (NF) mice developed AML within 4 weeks post-transplantation with significant leukocytosis (mean WBC 178.6 K/μL, mean Mono 43.9 K/μL), anemia (mean HCT 19.9%), and cKit+ peripheral blood cells (mean 11.5% of Cd45.2 cells; **Figure 4b, Extended Data Fig. 5b-c)**. The triple-mutant *Dnmt3a^Lox-R878H^*-*Npm1^Frt-c^*-*Flt3^GL-ITD^* (DNF) developed leukocytosis (mean WBC 104.25 K/μL, Mono 25.9 K/μL) and anemia (mean HCT 18.7%) consistent with AML, albeit with a slightly longer latency (DNF 6w vs. NF 4w). Single-mutant *Npm1^Frt-c^* (N) and double-mutant *Dnmt3a^Lox-R878H^*-*Npm1^Frt-c^* (DN) cells reliably engrafted but only developed leukemia at longer latency (median survival N 7.7w vs. DN 11.5w, log rank *p* ≤ 0.0082**; Figure 4b)**. We observed monoallelic *Dnmt3a^Lox-R878H^*-specific enrichment of LT-HSCs in addition to other genotype-specific alterations in downstream progenitor and mature cells (**Figure 4c, Extended Data Fig. 5d-e).** Specifically, while *Flt3^GL-ITD^* appeared to be the dominant driver of LSK expansion, antecedent *Dnmt3a^Lox-R878H^* mutations resulted in a further increase in this population (mean LSK% of Lin^-^ NF: 3.8% vs DNF: 9.52%, *p* ≤ 0.0038; **Figure 4d)**. These results indicate that while mutations in *Dnmt3a* promote pre-leukemic HSPC expansion, mutations in *Flt3* were the dominant drivers of immunophenotypic HSPC expansion at leukemic transformation.

To assess if the conserved clone-specific immunophenotypic properties we observed in our murine models were seen in clinical isolates from AML patients, we performed single-cell DNA-sequencing (scDNA-seq) with surface immunophenotyping on 97,086 cells across 4 patient samples possessing a combination of mutations in *DNMT3A, NPM1* and *FLT3* **(Extended Data Fig. 6a)**. We observed the expected decrease in CD34 expression in *NPM1* mutant clones (log2FC= −1.81, *FDR* ≤ 4.01×10^-206^), matching both the well-established clinical immunophenotype^23,24^ and our *Npm1^Frt-c^-Flt3^Frt-ITD^* model **(Extended Data Fig. 6b)**. Clustering based on immunophenotype revealed 6 communities of cells with representation of most clusters across the 4 samples (**Figure 4e, Extended Data Fig. 6c)**. We identified differentially expressed surface markers between the communities such that cells in community 1 expressed primitive HSPC markers including CD90 (92% of cells, log2FC 1.79), cells in community 2 cells expressed maturation markers such as CD14 (92% of cells, log2FC 1.46), and community 3 cells represented a progenitor enriched fraction with increased CD45RA expression (83% of cells, log2FC 1.36) (**Figure 4e, Extended Data 6c-d)**. We found clones were differentially distributed across the communities with *DNMT3A-FLT3* (DF) mutant clones most enriched in community 3 progenitor-like cells (66% of DF cells) and *NPM1-FLT3* (NF) cells enriched in community 2 differentiated cells (58.9%) (**Figure 4e-f)**. Finally, triple-mutant *DNMT3A-NPM1-FLT3* (DNF) cells were most enriched within the community 1 stem-like cells (49%), mirroring *DNF* mice which had the largest proportion of LSKs (**Figure 4d,f)**. These data confirm that different mutational combinations which promote leukemic transformation have differential impact on the leukemic hierarchy.

### Orthogonal chemical-genetic tools for modeling leukemia

A major limitation of the approaches described above and with other studies exploring mutational activation/cooperativity is the need for transplantation/adoptive transfer, which represents a “bottleneck” that selects out specific clones. To enable sequential mutational induction in the absence of transplantation, we employed chemical-genetic approaches capable of inducing Flp and Dre activation which are orthogonal to existing TAM-inducible ER fusions. For Flp, we developed a trimethoprim (TMP)-stabilizing Flpo dihydrofolate reductase (FlpoDHFR) fusion^26^. In a FlpoDHFR encoding retroviral infection-transplantation model, *in vivo* TMP treatment induced aggressive AML in *Npm1^Frt-c^*-*Flt3*^Frt-ITD^ cells with the same latency and immunophenotypic characteristics as the TAM-inducible FlpoERT2 system **(Extended Data 7a).**

For Dre, we used a Stabilized Peptide Linkage (StaPL) approach ^27^ by splitting Dre with an HCV-derived NS3 protease resulting in constitutive autoproteolysis of the construct (**Figure 5a)**. Inhibition of proteolytic activity with the NS3 inhibitor asunaprevir (ASV) resulted in efficient recombination (mean±s.e.m. 91±0.79%), albeit with detectable recombinase activity in the absence of ligand (mean±s.e.m. 9.9±0.96; **Extended Data Fig. 7b)**. This background recombination was nearly eliminated by cloning an ornithine-decarboxylase protected degron^28^ adjacent to the NS3 cut site (DreStaPL-ODC) resulting in degradation of the N-terminal fragment of Dre (mean±s.e.m. 0.043±0.008; **Extended Data Fig. 7b)**. Critically, the DreStaPL-ODC was orthogonally inducible to both TAM-inducible CreER and the trimethoprim (TMP)-stabilizing Flpo dihydrofolate reductase (FlpoDHFR) fusions (**Figure 5b)**. Thus, to complement existing Flpo and Cre tools, we generated two new DreStaPL-ODC mouse lines targeting either the 3’ UTR of the endogenous *Vav1* locus or the *H11* safe harbor under the control of a synthetic CAG promoter **(Extended Data Fig. 7c)**. CAG:DreStaPL-ODC mice displayed robust recombinase inducibility *in vivo* with NS3 inhibition by grazoprevir (GZV) (mean±s.e.m. 24.83±3.28) whereas the Vav1:DreStaPL-ODC mice demonstrated reduced recombination activity (mean±s.e.m. 0.176±0.05; *p* ≤ 0.0001; **Extended Data Fig. 7d)**.

**Figure 5:**
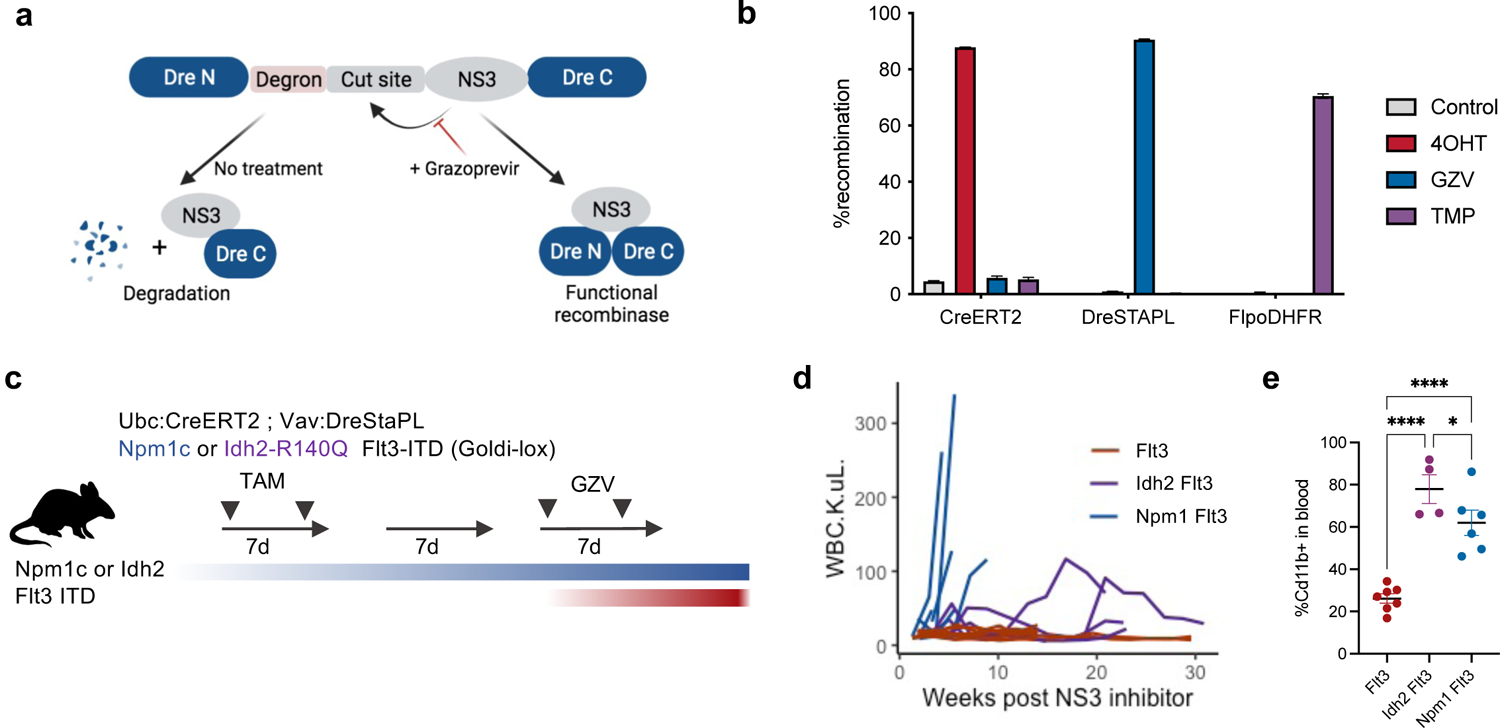
Chemically-inducible orthogonal recombinase activation. **a,** Schematic depicting DreSTAPL-ODC construct indicating that treatment with NS3 protease inhibitor results in complex formation of the split N and C-terminus of Dre. In the absence of NS3-inhibitin, proteolytic cleavage reveals the ODC degron leading to degradation of the N-terminus. **b,** Lin^-^ bone marrow from Rosa26: TdTomato (Lox-stop-Lox, Ai14), Rosa26:RLTG (Rox-stop-rox) or Rosa26:FLTG (Frt-stop-Frt) mice were infected with retroviral vectors encoding GFP and CreER, DreSTAPL-ODC, or FlpoDHFR respectively. Cultures were treated with the DMSO, 4-OHT (400nM), GZV (10μM) or TMP (1μM) as indicated. Barplot depicts recombination in infected population (%TdTomato+ of GFP+ cells). (n=3 per group). **c,** Experimental schema for primary mice with Ubc:CreERT2, Vav1:DreSTAPL-ODC, *Flt3^GL-ITD^* as well as either *Idh2^Lox-R140Q^* or *Npm1^Lox-C^.* Schema depicts dosing schedule of either TAM or GZV to activate Cre and Dre respectively. **d,** Line chart depicting in peripheral blood monocyte (K/μL) counts in *Flt3^GL-ITD^* (red), *Npm1^Lox-C^ Flt3^GL-ITD^* or *Idh2^Lox-R140Q^ Flt3^GL-ITD^* mice following GZV treatment. (n=4-13 per group). **e,** Stripchart indicating %Cd11b+ cells in the peripheral blood of mice depicted in (c-d) at either 16 weeks post GZV treatment (*Flt3^GL-ITD^* and *Idh2^Lox-R140Q^ Flt3^GL-ITD^*) or when symptomatic (*Npm1^Lox-C^ Flt3^GL-ITD^*). (n=4-7 per group). Error bars reflect the mean ± s.e.m.; and p-values are calculated by Fisher’s LSD test (e) *\* p ≤ 0.05* **** *p ≤ 0.0001*.

Finally, we sought to eliminate any reliance upon transplant for disease development. We proceeded with the Vav1:DreStaPL-ODC allele to model *Flt3^GL-ITD^* acquisition as a late, subclonal event. We crossed these alleles to mice harboring Ubc:CreER and either Cre-inducible *Npm1^Lox-c^* or *Idh2^Lox-R140Q^* mutations. Mice were administered TAM to activate either *Npm1^Lox-c^* or *Idh2^Lox-R140Q^* and then 2 weeks later treated with GZV to induce the *Flt3^GL-ITD^* allele (**Figure 5c)**. Despite the reduced *Flt3^GL-ITD^* induction frequency of Vav1:DreStaPL-ODC, *Npm1^Lox-c^-Flt3^GL-ITD^* mice developed an aggressive AML by 6 weeks post-GZV induction highlighting the potency of this mutational combination (**Figure 5d)**. In contrast, *Idh2^Lox-R140Q^*-*Flt3^GL-ITD^* (IF) mice developed a longer latency AML with more modest leukocytosis (mean WBC K/μL: IF 38.12 vs. NF 143.9, *p* ≤ 0.042) and reduced fraction of peripheral blood cKit+ cells compared to the *Npm1^Lox-c^*-*Flt3^GL-ITD^* model (mean±s.e.m.; IF 1.53±0.46 vs. NF 12.7±2.8, *p* ≤ 0.0042; **Figure 5d; Extended Data Fig. 7e)**. Despite the differences in disease latency, both models showed increased myeloid lineage commitment (mean Cd11b^+^% *Flt3*: 26.16%, IF: 77.95%, NF: 62.03%) and capacity to transplant AML into secondary recipients (**Figure 5e; Extended Data Fig. 7f-g)**. In sum, these chemical-genetic tools allow for modular, deterministic control of clonal evolution in malignancy in the absence of transplantation.

### Evaluating oncogene dependency of mutant-*Flt3*

We next sought to perturb disease evolution by abrogating the expression of mutant *Flt3 in vivo* with Cre-inducible mutant Flt3 reversion using the GOLDI-lox allele*. In vitro,* 4-OHT treatment led to reversal of the *Flt3^GL-ITD^* allele with concomitant reduction in cell number from leukemic *Npm1^Lox-c^*-*Flt3^GL-ITD^* (NF) mice but not *FlpoERT2 Npm1^Frt-c^ Flt3*^Frt-ITD^ mice **(Extended Data Fig. 8a-b)**. This decrease in cellularity was due in part to increased apoptosis as evidenced by increased AnnexinV+DAPI+ cells (*p* ≤ 0.0163; **Extended Data Fig. 8c)**. Cells from leukemic *Npm1^Lox-c^*-*Flt3^GL-ITD^* (NF) and *Idh2^Lox-R140Q^*-*Flt3^GL-ITD^* (IF) mice were then transplanted into lethally irradiated recipients and monitored for disease development at which point they were treated with TAM to delete the *Flt3^GL-ITD^* allele. *Flt3^GL-ITD^* ablation with TAM-mediated Cre expression was associated with a near complete reduction of WBC and decrease in spleen mass by 7 days in both the *NF* and *IF* mice (Pre-vs. Post-TAM WBC K/μL mean: NF 77.8 vs. 2.42; IF 106.7 vs. 0.82**; Figure 6a-b)**. Pathological analysis of sectioned bone marrow revealed a decrease in cellularity and nuclear condensation consistent with differentiation in both models (**Figure 6c)**. *Flt3^GL-ITD^* reversal also induced a reduction in lineage negative HSPCs in both models (NF: *p ≤* 0.0021, IF: *p ≤* 0.0006; **Figure 6d**), with NF mice expanding granulocytes and a Cd11c^+^ dendritic-like population upon TAM-treatment **(Extended Data 8d)**. While the IF mice displayed a reduction in LSKs (mean %LSK of Cd45.2 Control: 4.01% vs. TAM: 0.81%), NF mice had an increase in immunophenotypic LSKs (mean %LSK of Cd45.2 Control: 1.74% vs. TAM: 3.37%) with concomitant reduction in Lin^-^Kit^+^Sca1^-^ progenitor cells (**Figure 6d)**. Similar results were observed after 4-OHT administration in *ex vivo* culture of NF cells (*p* ≤ 0.0148**; Extended Data Fig. 8e)**.

**Figure 6:**
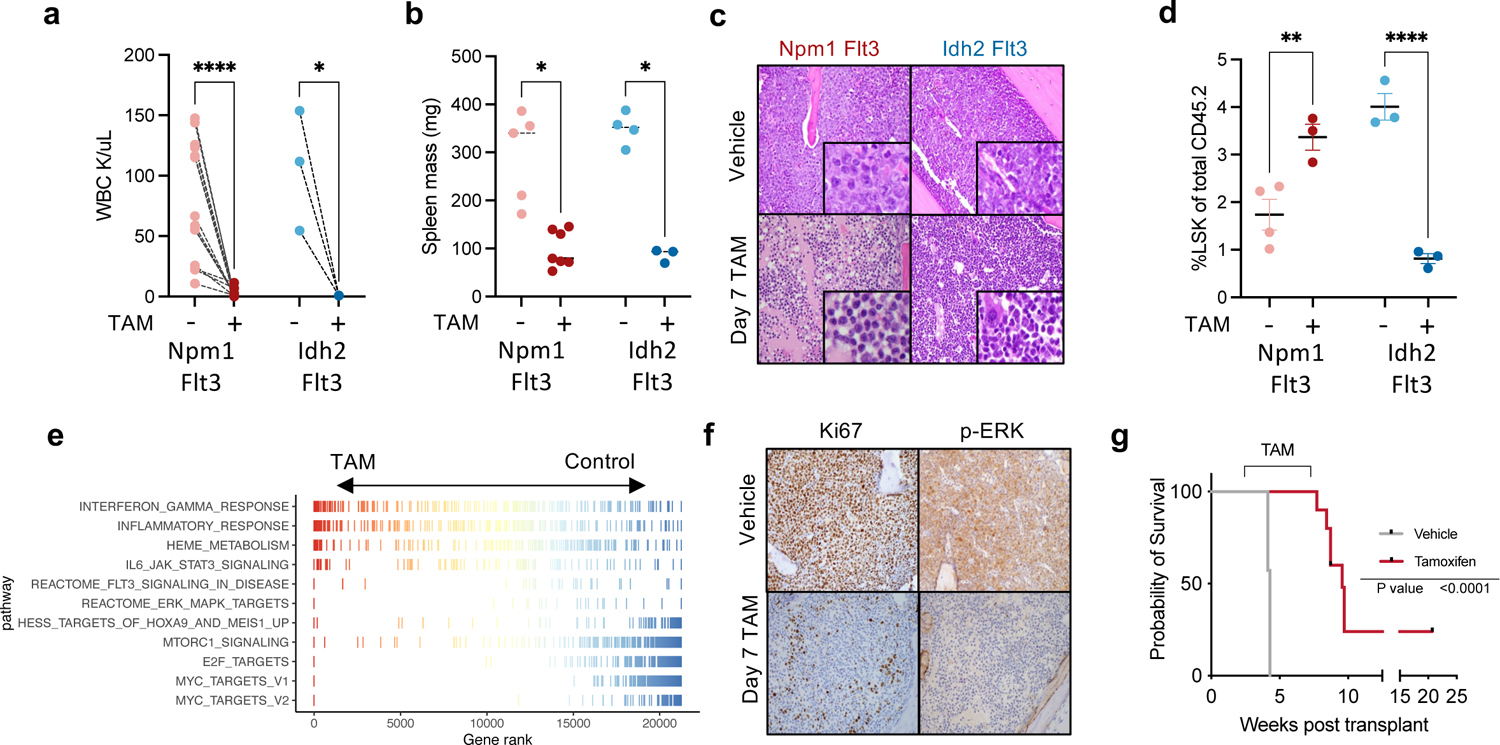
Reversible *Flt3* mutagenesis in acute myeloid leukemia Secondary transplant of *Npm1^Lox-C^ Flt3^GL-ITD^* (red; 20,000 cells) or *Idh2^Lox-R140Q^ Flt3^GL-ITD^* (blue; 500,000) from Figure 5 were engrafted into lethally irradiated secondary recipients with 500,000 Cd45.1 WT cells. After transplant mice were monitored for disease development (*Npm1^Lox-C^ Flt3^GL-ITD^* 4 weeks, or *Idh2^Lox-R140Q^ Flt3^GL-ITD^* 10 weeks), and then euthanized following 7 days of TAM treatment (a-g). **a,** Peripheral blood WBC (K/μL) before and after TAM treatment.(n=3-14 per group) **b,** Spleen mass in control and TAM treated mice. (n=3-7 per group). **c,** Bone marrow H&E stained from either control mice or after 7 days of TAM (400x). **d,** Dot chart depicting %LSKs of Cd45.2 cells in bone marrow from control and TAM treated mice for the indicated disease models. (n=3-4 per group). **e,** RNA-sequencing data from lin^-^ HSPCs purified from 7-day TAM treated *Npm1^Lox-C^ Flt3^GL-ITD^* or control mice. Gene ranks from GSEA are depicted as a linechart for the indicated HALLMARK gene set. **f,** Immunohistochemical staining on bone marrow for Ki67 (left) and phopsho-ERK1/2 (right) for untreated (top) and 7 day TAM treated (bottom) *Npm1^Lox-C^ Flt3^GL-ITD^* mice. **g,** Kaplan-Meier survival curve of mice engrafted with 500,000 Cd45.1 WT cells and 40,000 bone marrow cells from symptomatic *Npm1^Lox-C^ Flt3^GL-ITD^* mice from Figure 5. Mice were either treated with control chow (grey) or placed on a TAM-chow diet (red) for 4 weeks. (n=7-10) Error bars reflect the mean ± s.e.m.; and p-values are calculated by Dunn’s test (a,b), Fisher’s LSD test (d) and Log-Rank test (g). *\* p ≤ 0.05 ** p ≤ 0.01* *** *p ≤ 0.001* **** *p ≤ 0.0001*

This increase in phenotypic LSKs suggested that reversal of *Flt3*^ITD^ expression induces a compensatory increase in inflammation and resultant upregulation of *Sca1* expression. Gene expression profiling of *Npm1^Lox-c^*-*Flt3^GL-ITD^* HSPCs showed that TAM treatment was associated with an increase in inflammatory gene set primarily driven by an IFNγ response (GSEA NES: 2.39, *FDR* ≤ 8.8×10^-12^) and STAT3/IL6 signaling (GSEA NES: 1.39, *FDR* ≤ 3.3×10^-2^; **Figure 6e; Supplementary Table 1d)**. Consistent with this, measurement of serum cytokine levels showed an increase in IL6 (*p* ≤ 0.0011) levels **(Extended Data 8f**). We also observed a decrease in Hoxa9 and Meis1 target gene expression (NES −2.91, *FDR* ≤ 2.55×10^-19^) underscored by decreased expression of the transcription factors themselves (**Figure 6e; Extended Data Fig. 8g; Supplementary Table 1e)**. Additionally, we observed a reduction in gene expression signatures for cell cycle related E2F targets, MYC targets and ERK pathway engagement (**Figure 6e; Extended Data Fig. 8g)**, with immunohistochemical analysis supporting these findings (**Figure 6f, Extended Data 8h)**. Finally, *Npm1^Lox-c^*-*Flt3^GL-ITD^* HSPCs showed re-engagement with erythropoiesis by an increase in genes associated with Heme synthesis (NES 1.92, *FDR* ≤ 8.5×10^-7^) as well as increased expression of *Gata1* (*FDR* ≤ 2.56×10^-7^) and *Epor* (*FDR* ≤ 8.9×10^-17^; **Figure 6e; Extended Data Fig. 8g)**. Collectively, these alterations indicate that mutant-*Flt3* deletion in AML cells is associated with a reduction in proliferation, reduced expression of key hematopoietic self-renewal transcriptional regulators, and attenuated signal transduction.

We observed an initial decrease in mutant chimerism, and long-term treatment with TAM resulted in an increase in overall survival, with a subset of *Npm1^Lox-c^*-*Flt3^GL-ITD^* (2/10) showing no evidence of disease relapse (**Figure 6g; Extended Data Fig. 8i)**. In mice with recurrent AML, we observed AMLs with residual Dre-activated *Flt3^GL-ITD^* that was not deleted by Cre as well as evidence of continued Cre-mediated *Flt3^GL-ITD^* deletion **(Extended Data Fig. 8j)**. These findings suggest that reversal of mutant *Flt3^ITD^* can reduce leukemic burden and improve survival, although leukemic clones can recur through both Flt3^ITD^-dependent and independent mechanisms.

## Conclusion

Advances in whole genome sequencing and single-cell genomics have offered refined resolution into the evolutionary processes underlying cancer development and response to therapy^1-3,29,30 31^. While GEMMs have served as a robust means to model disease, spatio-temporal control of stepwise somatic alterations has remained elusive. Here we show that the use of orthogonal recombinases allows one to model sequential mutational acquisition and to develop AML models which follow the trajectories observed in patient samples. This led to insights into leukemic transformation and will inform studies of how mutational order, cell type of mutation acquisition, and mutational clonal representation impact leukemic transformation and the response to therapy.

Most importantly, these studies provide a template by which multiple mutations can be deterministically and sequentially induced *in vivo*, without the selection bottlenecks induced by transplantation,^32^ simultaneous CRISPR editing of different mutant alleles,^17,18^ or ectopic mutant gene expression^32 33,34^. Each of these approaches will have an important role in modeling cancer evolution and clonal complexity, and there may be advantages to combining endogenous targeting and multiplex CRISPR editing to better model mutational complexity *in vivo*. We believe that the use of orthogonal recombinases will allow investigators to induce different mutations in different susceptible populations and to use single-cell technologies to both follow clonal evolution over time and delineate how mutational events coordinately alter cell state. Moreover, the use of reversible systems to activate and inactivate mutant alleles from their endogenous loci builds upon previous transgenic/reversible shRNA models^35-37^ to allow more precise, mutant-specific reversible gene activation. This allows for new insights into oncogenic dependency and into how cancer disease alleles alter the transcriptional, epigenetic and phenotypic output of cancer cells. Moreover, this system can be used to credential new mutant-specific dependencies in different malignant contexts and to delineate mechanisms of oncogenic dependency and escape which can inform preclinical and clinical therapeutic studies.

There are many limitations to our current approach, as we are limited to simultaneously modeling a small number of disease alleles, and most human cancers present as an amalgam of clonal complexity which changes throughout disease progression and in response to specific therapies^38^. Moreover, our system does not account for the important role of non-genetic factors, including epigenetic remodeling, in cancer evolution and therapeutic response,^6^ and we have not investigated how complex mutant clones interact with the local and system tumor microenvironment and immune system. However, our work provides a roadmap to investigate clonal evolution and inter-clonal interactions in different malignant contexts, including in the context of cell non-autonomous factors which alter mutational rate and/or selection. As our ability to fill in the atlas of somatic genetic events which occur from normal cells to pre-malignant somatic expansion to overt transformation in the human context expands,^39^ the continued development of more advanced systems to model clonal evolution will afford an unprecedented view of the mechanisms mediating transformation with biologic, therapeutic, and translational import.

**Extended Data Fig. 1:**
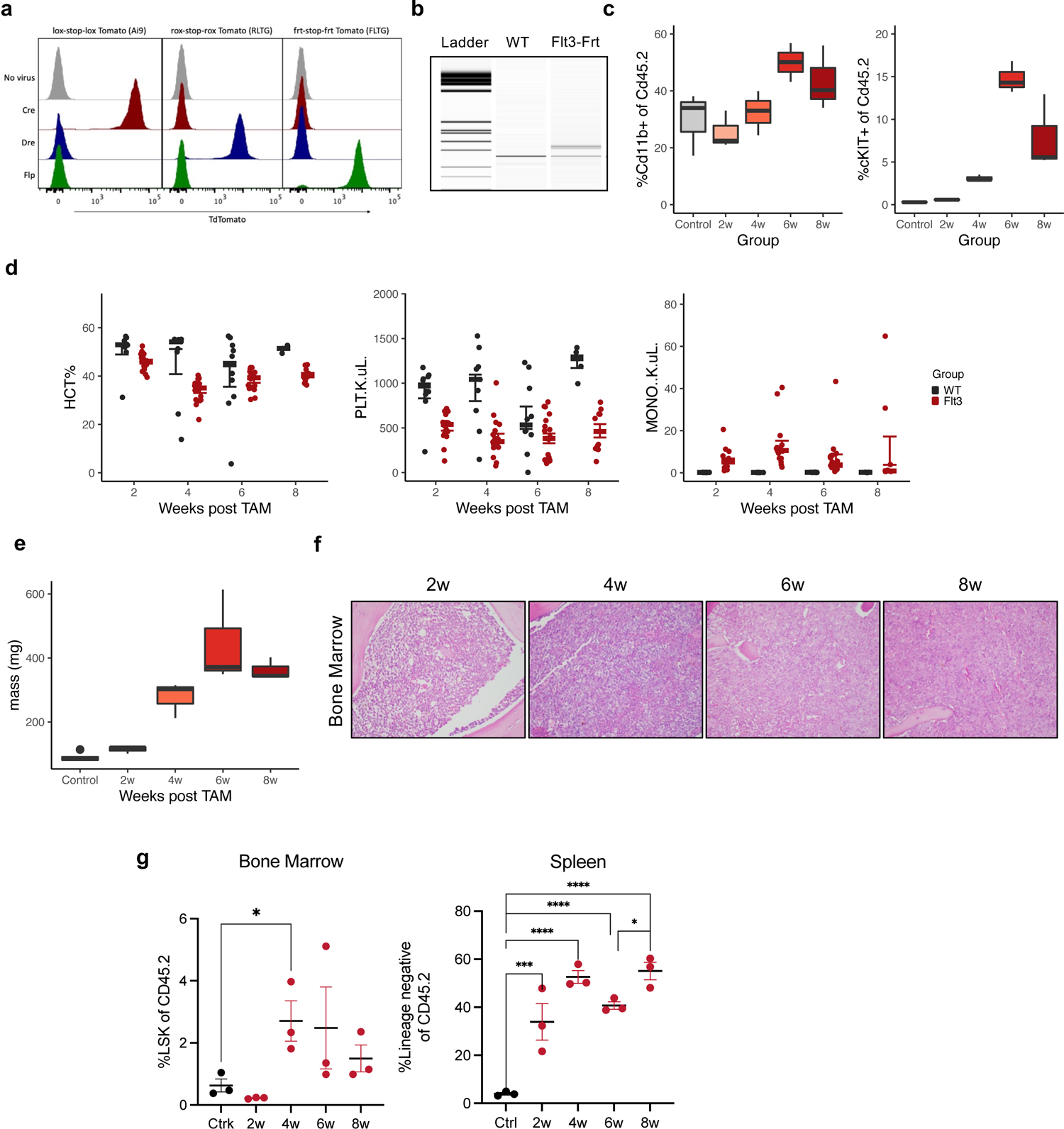
Flp-inducible *Flt3^Frt-ITD^* mice demonstrate aberrant hematopoietic phenotypes. **a,** Lin^-^ bone marrow was isolated from mice encoding a transcriptional stop cassette flanked by either Lox, Rox or Frt sites. Cells were infected with an GFP+ retrovirus encoding either Cre, Dre or Flp. Excision of the stop cassette in all three reporter lines results in TdTomato expression. Histogram depicts TdTomato expression in GFP+, virally infected cells. **b,** Rosa26:FlpoERT2 *Flt3^Frt-ITD^* or WT mice were treated with TAM, and DNA was isolated for PCR to evaluate excision at the Flt3 locus. Lower band indicates WT *Flt3* upper band indicates Flp-mediated inversion. PCR was visualized with capillary electrophoresis (Qiaxcel). **c,** Lethally irradiated Cd45.1 recipient mice were transplanted with Rosa26:FlpoERT2 *Flt3^Frt-ITD^* bone marrow and treated with TAM 4 weeks after engraftment. Boxplot depicting total myeloid (Cd11b^+^, left) and cKIT+ (right) cells of Cd45.2 cells in bone marrow transplant at 2, 4, 6 and 8 weeks post TAM. Control mice are *Flt3^WT^* and 4 weeks post TAM. (n=3 per group) **d,** Hematocrit (%, left), platelets (K/μL, middle) and monocytes (K/μL, right) from primary Rosa26:FlpoERT2 *Flt3^Frt-ITD^* or WT at 2, 4, 6, and 8 weeks post TAM. (n=10-19 per group) **e,** As in (c), mice were transplanted with Rosa26:FlpoERT2 *Flt3^Frt-ITD^* bone marrow and treated with TAM 4 weeks after engraftment. Boxplots depict spleen mass 2, 4, 6 and 8 weeks post TAM. Control mice are *Flt3^WT^* and 4 weeks post TAM. **f,** H&E staining of bone marrow from mice described in panels c and e (400X). (n=3 per group) **g,** Dot chart depicting %LSK of Cd45.2 cells in bone marrow (left) or %Lin^-^ of Cd45.2 in spleen (right) from mice transplanted with Rosa26:FlpoERT2 *Flt3^Frt-ITD^* at 2, 4, 6, and 8 weeks post TAM. (n=3 per group). Error bars reflect the mean ± s.e.m.; and p-values are calculated by Fisher’s LSD test (g), *\* p ≤ 0.05* *** *p ≤ 0.001* **** *p ≤ 0.0001.* Boxplots depict median and IQR with whiskers extending to 1.5*IQR.

**Extended Data Fig. 2:**
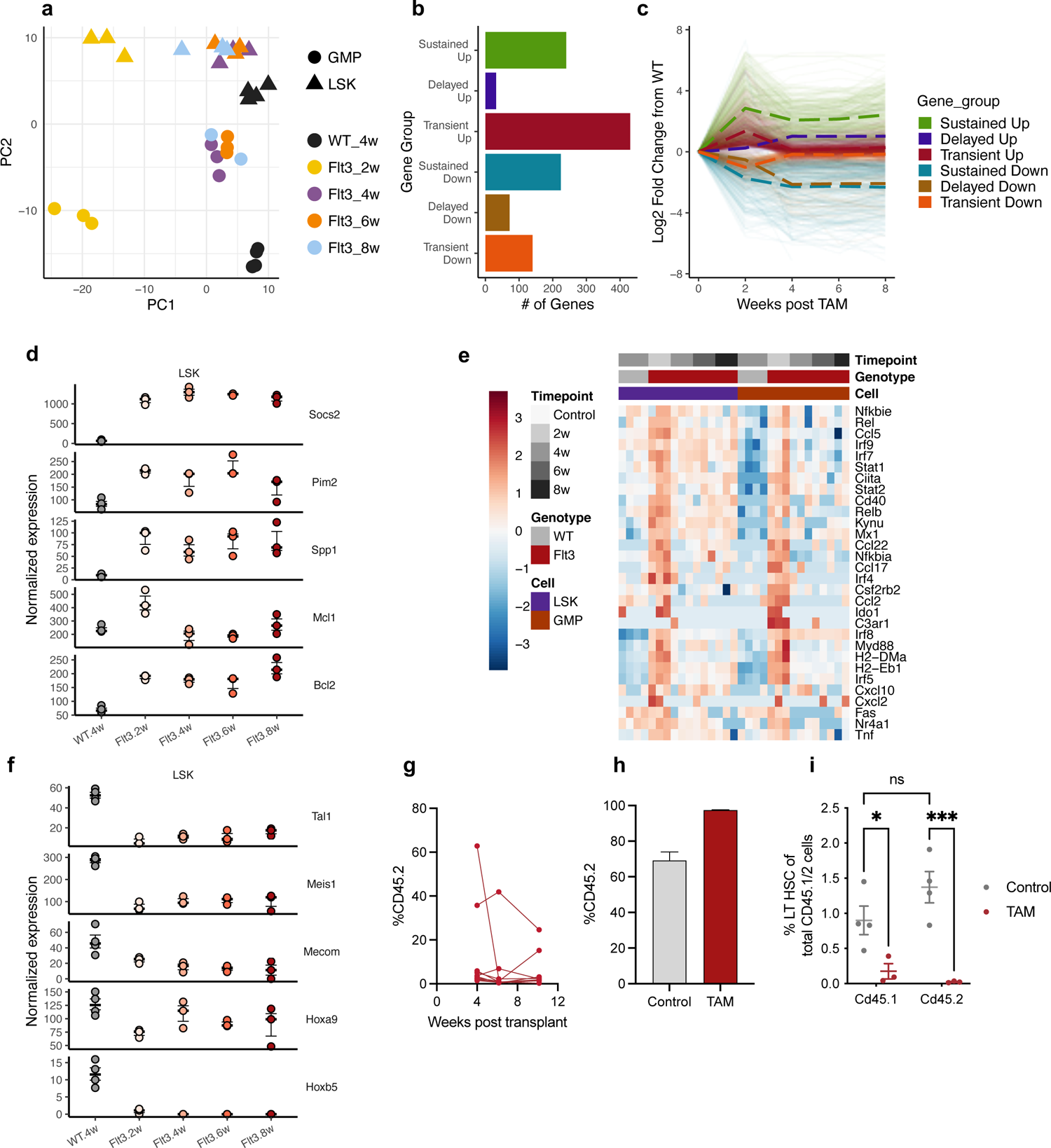
Transcriptional alterations associated with stem cell dysregulation in *Flt3^Frt-ITD^* mice. **a,** Principal component analysis on RNA-sequencing data of the 1000 most variable genes across LSKs and GMPs sorted from WT or *Flt3^Frt-ITD^* LSKs at either 2, 4, 6 or 8 weeks post TAM. (n=2-3 per group). **b,** Barplot depicting number of downregulated (blue) and upregulated (red) differentially expressed genes in LSKs for the indicated comparisons. **c,** Row-normalized heatmap depicting average expression levels for gene clusters identified by K-means clustering. **d,** Boxplot depicting normalized gene expression values in WT or *Flt3^Frt-ITD^* LSKs at either 2, 4, 6 or 8 weeks post TAM. (n=2-3 per group) **e,** Row-normalized heatmap of RNA-sequencing data in WT or *Flt3^Frt-ITD^* LSKs and GMPs at either 2, 4, 6 or 8 weeks post TAM. **f,** Boxplot depicting normalized gene expression values for stem related genes in WT or *Flt3^Frt-ITD^* LSKs as in (d). (n=2-3 per group) **g,** Primary Rosa26:FlpoERT2 *Flt3^Frt-ITD^* mice were treated with TAM at 6 weeks of age, and bone marrow was harvested 24 weeks after TAM. Cells (2×10^6^) from *Flt3^Frt-ITD^* mice were mixed with Cd45.1 whole bone marrow (3×10^5^) and transplanted into lethally irradiated Cd45.1 recipients. Linechart depicts Cd45.2 chimerism in peripheral blood. (n=10) **h,** Rosa26:FlpoERT2 *Flt3^Frt-ITD^* was mixed with Cd45.1 WT cells 1:1 and transplanted into lethally irradiated Cd45.1 recipient mice. Mice were treated with TAM 4 weeks post engraftment, and euthanized 16 weeks post TAM. Barplot depicts Cd45.2 chimerism at 16 weeks post TAM. (n=3-4 per group). **i,** As in (h), whole bone marrow was isolated from mice 16 weeks post TAM, and LT-HSC (lin^-^ cKIT^+^ Sca1^+^ Cd48^—^ Cd150^+^) were quantified by flow cytometry in both the Cd45.1 WT and Cd45.2 mutant compartment. (n=3-4 per group). Error bars reflect the mean ± s.e.m.; and p-values are calculated by Fisher’s LSD test (i), *\* p ≤ 0.05* *** *p ≤ 0.001*.

**Extended Data Fig. 3:**
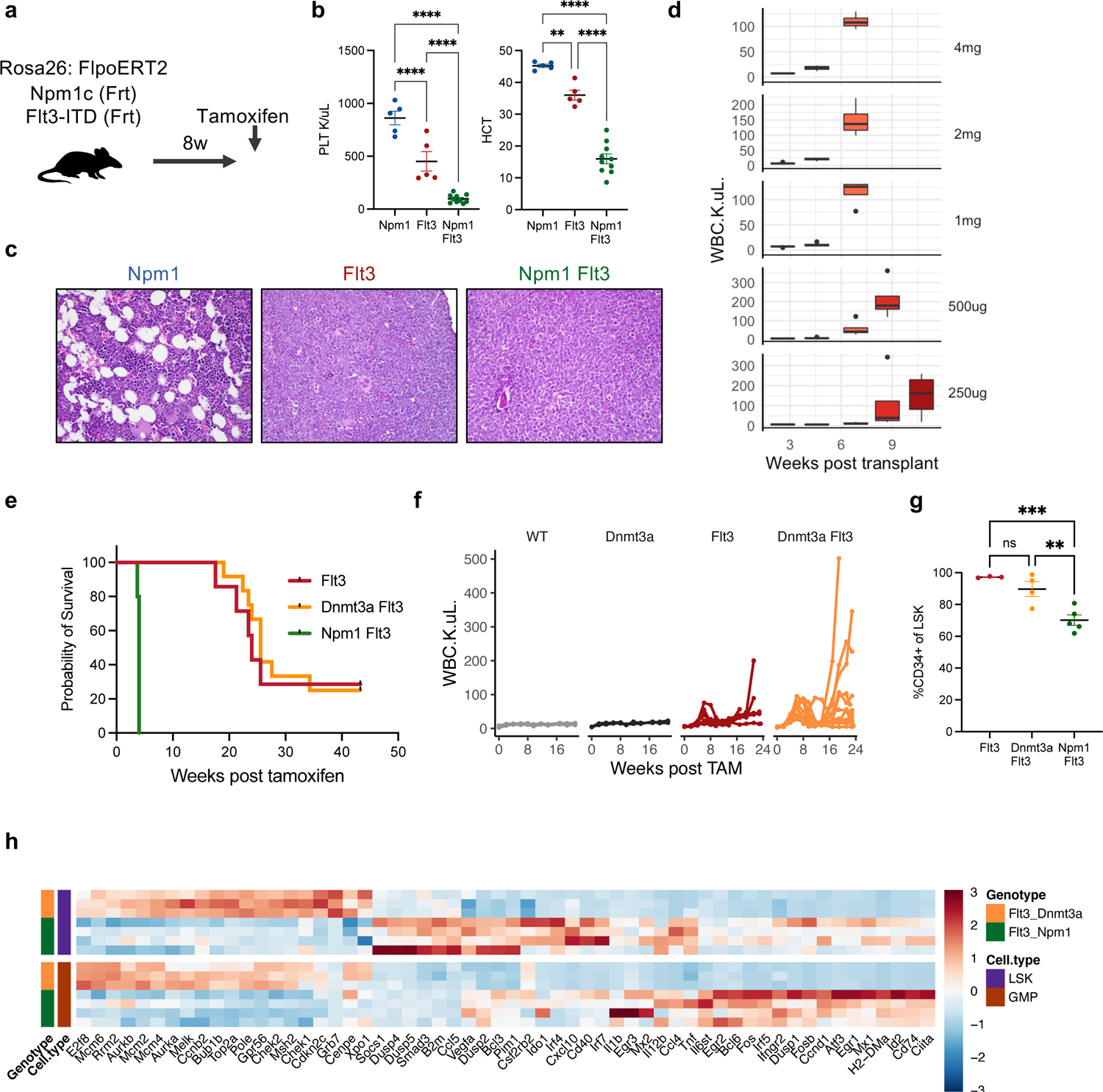
*Npm1*^Frt-C^ and *Dnmt3a^Lox-R878H^* license *Flt3^Frt-ITD^* for transformation. **a,** Schematic indicating simultaneous activation of *Npm1*^Frt-C^ and *Flt3^Frt-ITD^* by TAM-mediated FlpoERT2 activation. **b,** As in Figure 2b, bone marrow was transplanted into lethally irradiated recipient mice. After transplant (4w) mice were treated with TAM to activate the indicated genotypes. Stripchart indicates peripheral blood platelets (K/μL) (left) or hematocrit (%) (right) 4 weeks post TAM. (n=5-10). **c,** H&E staining on bone marrow from the indicated genotypes 4 weeks post TAM as in (b). **d,** Bone marrow from Rosa26:FlpoERT2 *Npm1*^Frt-C^ *Flt3^Frt-ITD^* mice were transplanted into lethally irradiated recipients and treated with the indicated titration of TAM. Boxplots depict WBC (K/μL) at biweekly bleeds post TAM. (n=3 per group). **e,** Kaplan-Meier survival curve of *Flt3^Frt-ITD^* (red), *Dnmt3a^Lox-R878H^ Flt3^Frt-ITD^* (orange), and *Npm1^Lox-C^ Flt3^Frt-ITD^* mice following TAM-mediated Rosa26:FlpoERT2 activation. (n=5-12 per group). **f,** Line chart depicting monocytes (K/μL) of WT, *Dnmt3a^Lox-R878H^, Flt3^Frt-ITD^* and *Dnmt3a^Lox-R878H^ Flt3^Frt-ITD^* mice following TAM-mediated activation of Rosa26:FlpoERT2. (n=3-12 per group). **g,** Fraction of Cd34+ cells in LSKs in bone marrow from *Flt3^Frt-ITD^, Npm1^Frt-C^ Flt3^Frt-ITD^,* and *Dnmt3a^Lox-R878H^ Flt3^Frt-ITD^* mice. (n=3-5 per group). **h,** Column-normalized heatmap of gene expression data from LSKs (bottom, purple) or GMPs (top, orange) from *Npm1^c-Frt^ Flt3^Frt-ITD^* (green) or *Dnmt3a^Loox-R878H^ Flt3^Frt-ITD^* (orange) leukemic mice (rows) with each column depicting the indicated gene. Error bars reflect the mean ± s.e.m.; and p-values are calculated by Fisher’s LSD test (b,g), *\*\* p ≤ 0.01* *** *p ≤ 0.001* **** *p ≤ 0.0001.* Boxplots depict median and IQR with whiskers extending to 1.5*IQR.

**Extended Data Fig. 4:**
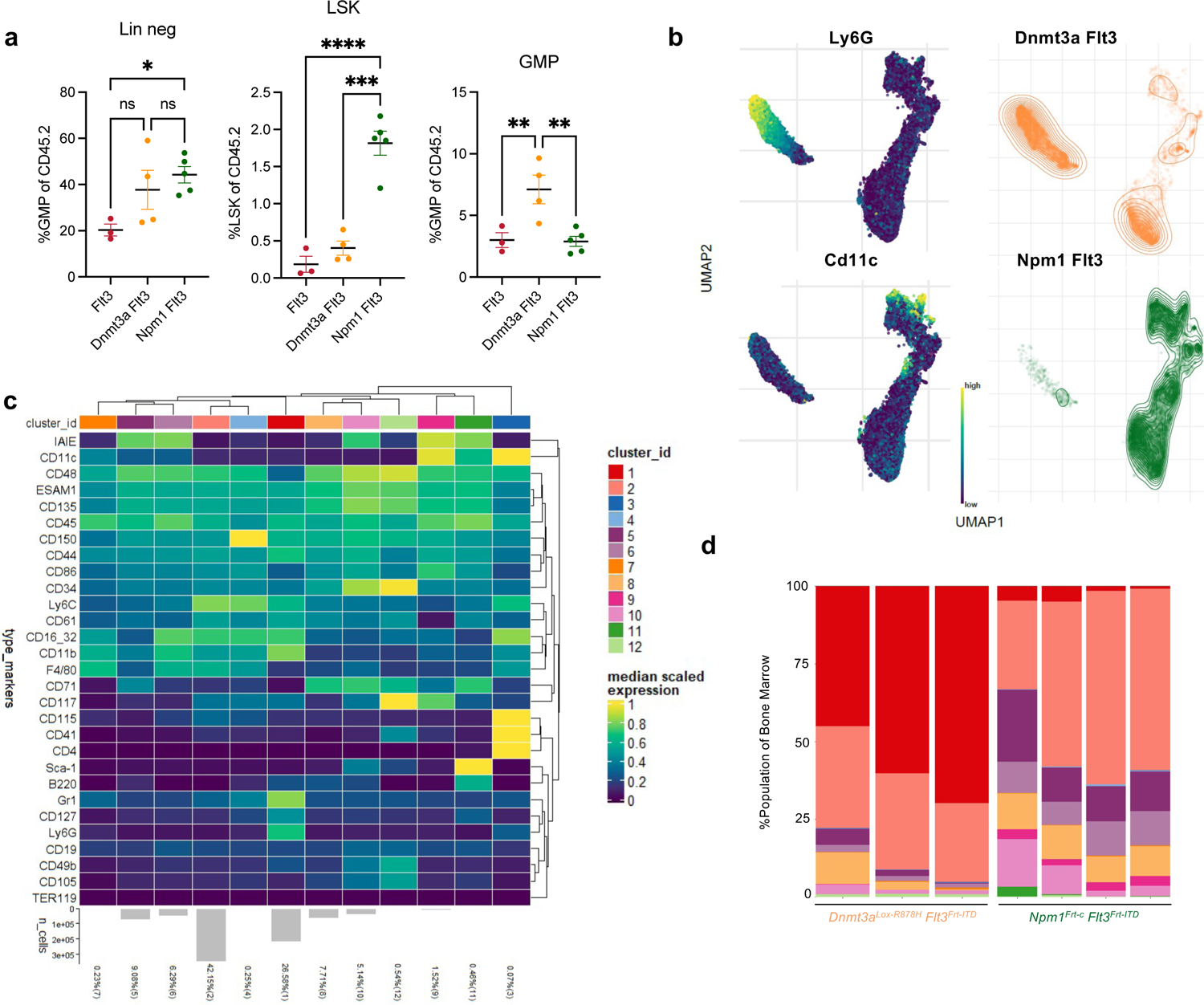
Immunophenotypic characterization of *Dnmt3a-Flt3* and *Npm1-Flt3* leukemias. **a,** Dot chart depicting %lin^-^ (left), %LSK^+^ (middle) and GMP (right) of Cd45.2 cells in bone marrow from *Flt3^Frt-ITD^, Npm1^Frt-C^ Flt3^Frt-ITD^,* and *Dnmt3a^Lox-R878H^ Flt3^Frt-ITD^* mice. (n=3-5 per group) **b,** UMAP depicting CyTOF-derived abundance levels of Ly6G (top left) and Cd11c (bottom left) in bone marrow from leukemic mice. Relative frequency of cells from *Dnmt3a^Lox-R878H^ Flt3^Frt-ITD^* (top right) and *Npm1^Frt-C^ Flt3^Frt-ITD^* (bottom right) are depicted as density contour plots. **c,** Heatmap indicating CyTOF-derived median scaled expression value for markers (rows) in each cluster of cells (columns). Cluster abundance is depicted as a barplot on the bottom of the graph. **d,** Stacked barplot depicting abundance of cells in clusters from (d) for *Dnmt3a^Lox-R878H^ Flt3^Frt-ITD^* (left) and *Npm1^Frt-C^ Flt3^Frt-ITD^* (right) leukemias. Error bars reflect the mean ± s.e.m.; and p-values are calculated by Fisher’s LSD test (a), *\* p ≤ 0.05 ** p ≤ 0.01* *** *p ≤ 0.001* **** *p ≤ 0.0001*.

**Extended Data Fig. 5:**
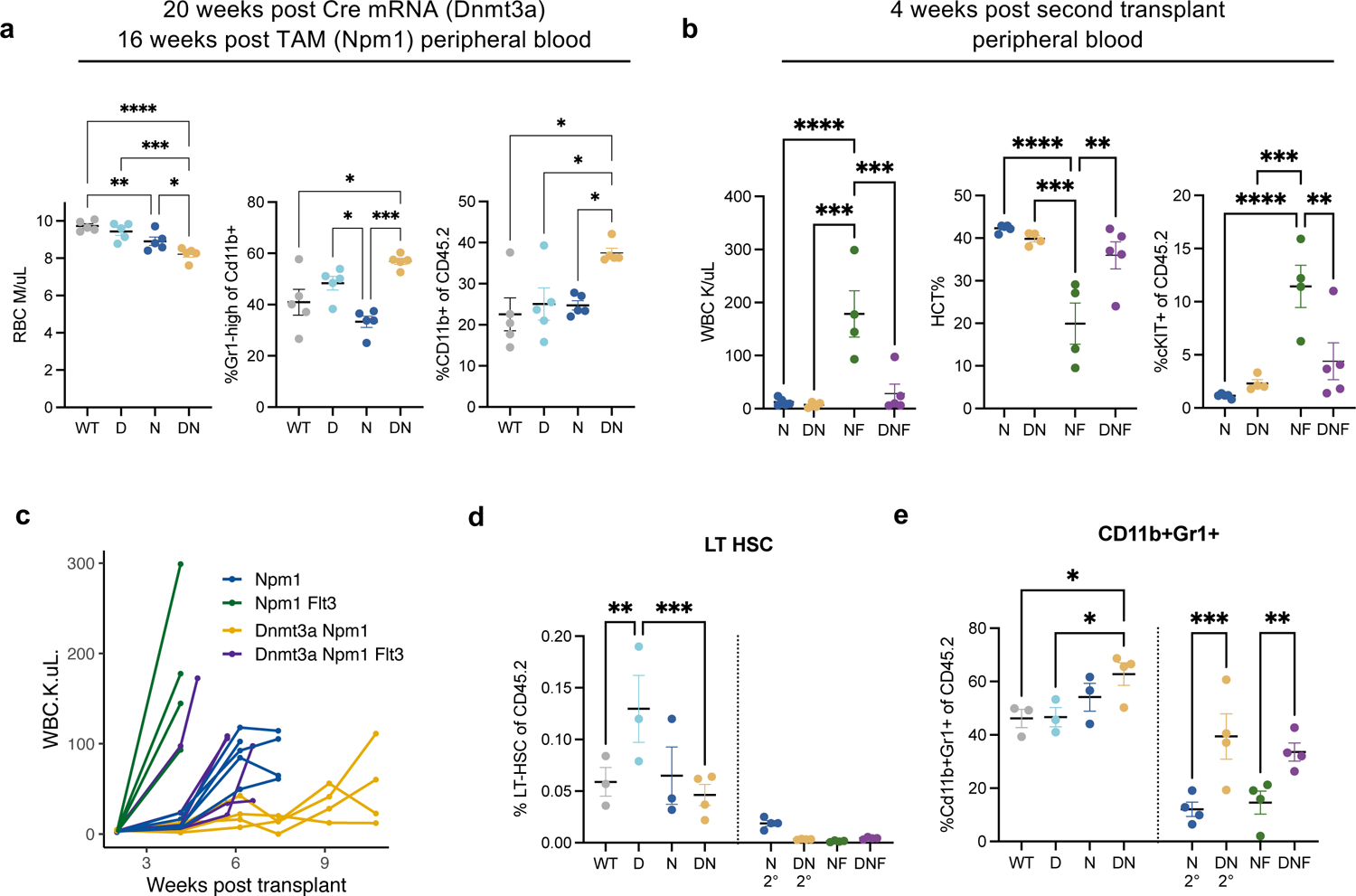
Immunophenotypic characterization of sequential mutagenesis models Related to experimental scheme depicted in Figure 4. **a,** Flow cytometric analysis in the peripheral blood at the end of the first transplant (20 weeks post Dnmt3a^Lox-R878H^/Cre, 16 weeks post *Npm1*^Frt-C^/Flp) for total myeloid cells (Left, %Cd11b+ of total Cd45.2), monocytic cells (middle, %Gr1^mid^ of Cd45.2), and granulocytic cells (right, %Gr1^high^ of Cd45.2). (n=5 per group). **b,** Flow cytometric analysis in the peripheral blood 4 weeks after the second transplant (4 weeks after *Flt3^GL-ITD^*/Dre or mock electroporation) for cKIT+ cell (Left), monocytic cells (middle, %Gr1^mid^ of Cd45.2), and granulocytic cells (right, %Gr1^high^ of Cd45.2). (n=4-5 per group). **c,** Peripheral blood WBC count (K/μL) in secondary transplant for the indicated genotypes. (n=5 per group). **d-e,** Strip chart from mice depicted in Figure 4c with y-axis indicating %LT-HSC (d), or %Cd11b^+^Gr1^+^ (e) cells within the Cd45.2 compartment for the indicated genotypes. N 2° and DN 2° indicate secondary transplant with mock mRNA electroporation. (n=3-4 per group). Error bars reflect the mean ± s.e.m.; and p-values are calculated by Fisher’s LSD test (a,b,d,e), *\* p ≤ 0.05 ** p ≤ 0.01* *** *p ≤ 0.001* **** *p ≤ 0.0001*.

**Extended Data Fig. 6:**
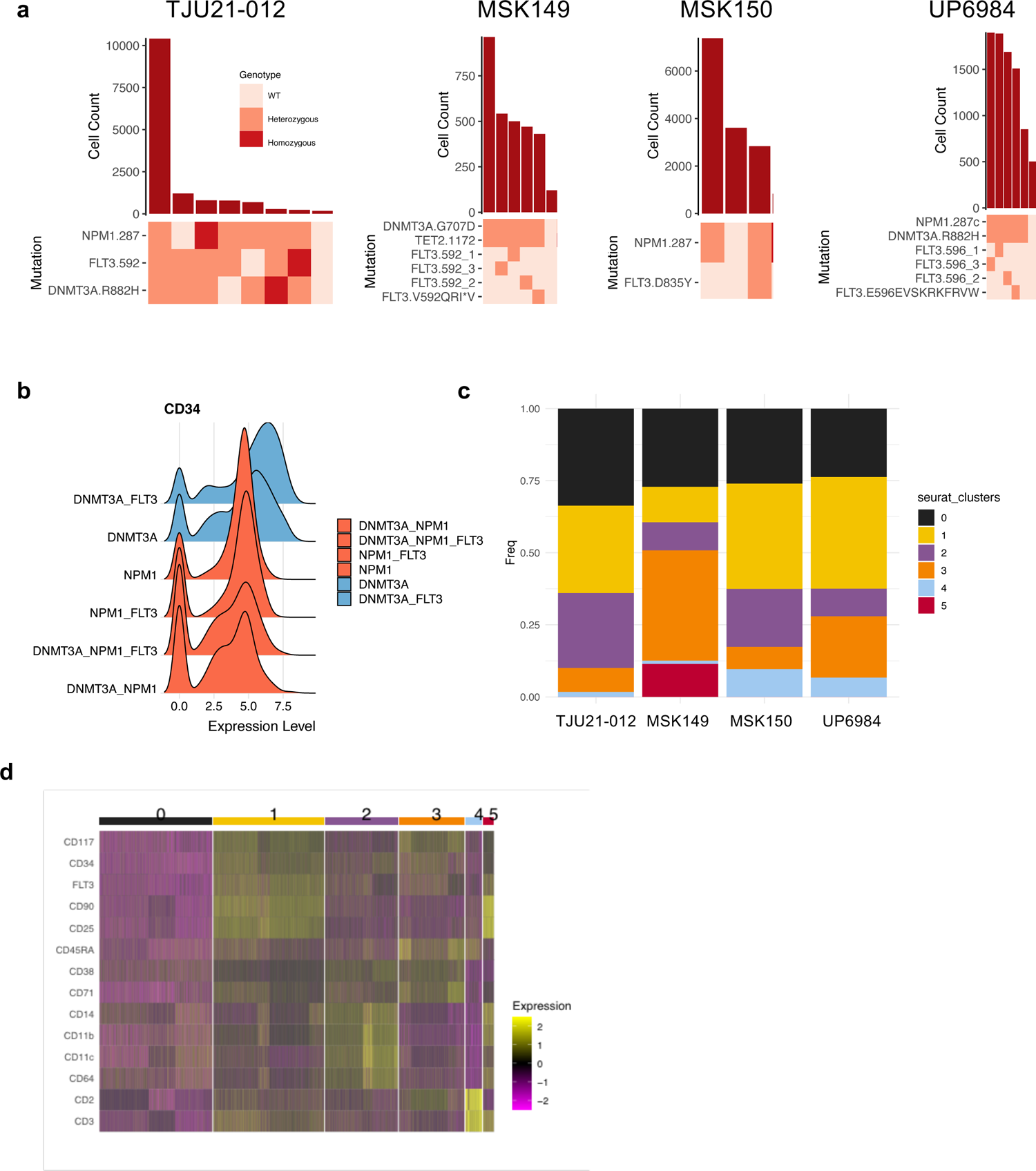
Single cell DNA sequencing and surface immunophenotyping of AML samples. **a,** Clonographs depicting mutation abundance for each sample. Top bar plot depicts number of cells identified with a given genotype and ranked by decreasing frequency. Bottom, heat map indicates mutation zygosity for each clone. **b,** Histogram depicting log-normalized CD34 expression in indicated clones, with *NPM1^mutant^* clones depicted in red and *NPM1^WT^* clones depicted in blue. **c,** Stacked barplot depicting relative abundance of different cell communities for each sample. **d,** Row normalized heatmap depicting expression of select markers across the different cell communities, high expression indicated in yellow and low expression indicated in purple.

**Extended Data Fig. 7:**
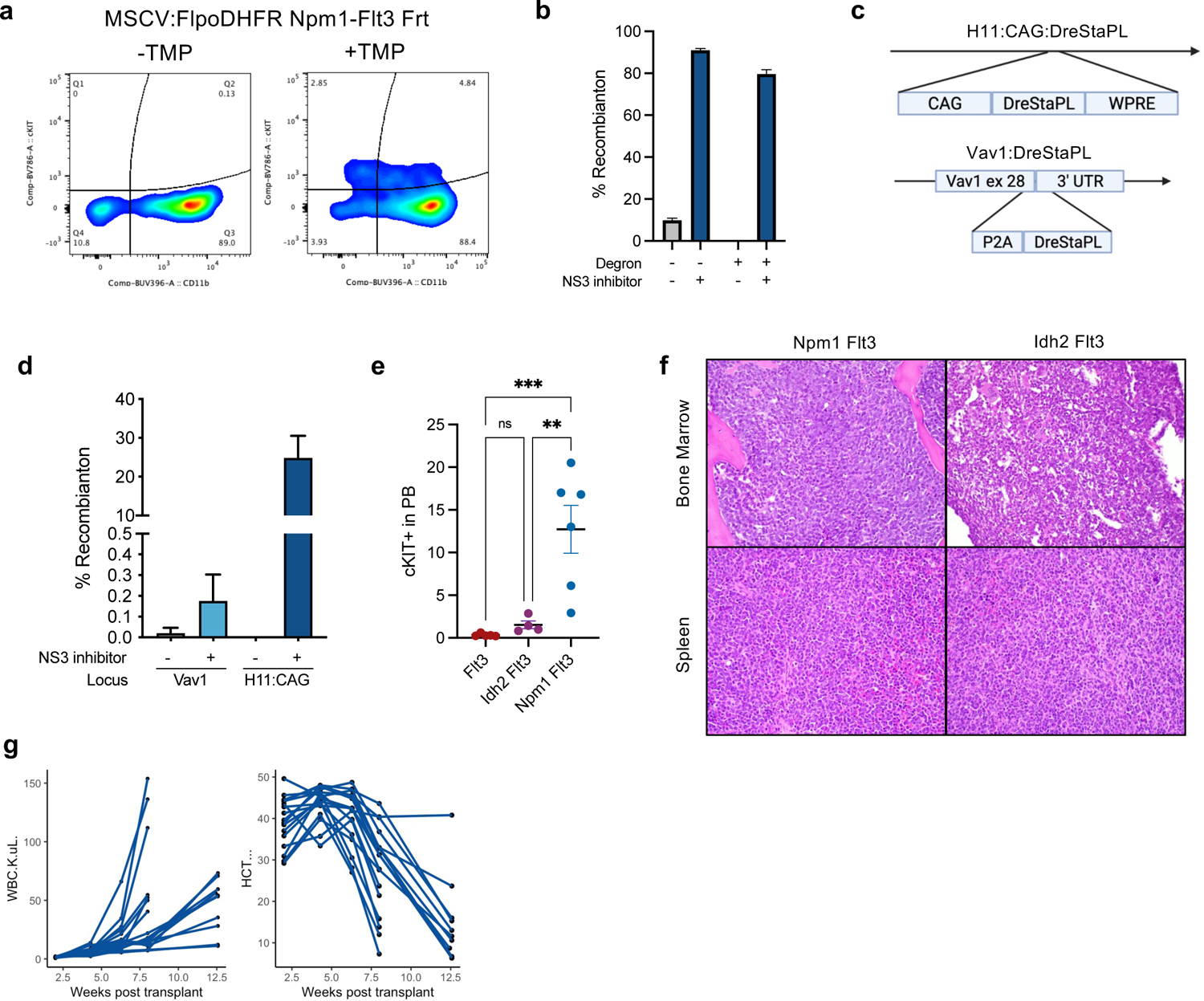
Inducible recombinases for modeling *Flt3* activation and leukemogenesis. **a,** Bone marrow was isolated 6 days after 5-FU treatment from *Npm1^Frt-C^ Flt3^Frt-ITD^* mice and infected with MSCV:FlpoDHFR IRES GFP virus. Cells were transplanted with 200,000 Cd45.1 WT cells into lethally irradiated recipients and engraftment was monitored by flow cytometry. Mice were treated with and without the FlpoDHFR stabilizing agent trimethoprim (TMP). Representative flow plot indicating accumulation of cKIT+ cells in the peripheral blood following TMP-treatment. **b,** Lin^-^ bone marrow from Rosa26:RLTG mice were infected with MSCV:DreSTAPL(ODC)-IRES-GFP virus with and without the ODC degron and treated with 10uM asunaprevir (ASV). Barplot depicting recombination (%TdTomato+ from Rosa26:RLTG). (n=3 per group). **c,** Schematic representation of CAG:DreSTAPL-ODC WPRE knockin into the H11 locus (top) or P2A-DreSTAPL-ODC knockin into the terminal exon of the Vav1 locus (bottom) replacing the endogenous stop codon. **d,** DreSTAPL-ODC knockin mice from (d) were crossed to the Rosa26:RLTG reporter and treated with GZV. Recombination was assessed by %TdTomato+ in the peripheral blood 2 weeks post GZV treatment. (n=3-5 per group). **e,** Stripchart depicting cKIT+ cells in the peripheral blood of *Flt3^GL-ITD^, Idh2^Lox-R140Q^ Flt3^GL-ITD^* or *Npm1^Lox-C^ Flt3^GL-ITD^* mice 12 weeks after GZV treatment (or at symptomatic endpoint for *Npm1^Lox-C^ Flt3^GL-ITD^* mice). (n=4-6 per group). **f,** H&E staining in *Npm1^Lox-C^ Flt3^GL-ITD^* (left) and *Idh2^Lox-R140Q^ Flt3^GL-ITD^* (right) for bone marrow (top) and spleen (bottom) from symptomatic mice (400X) .**g,** Whole bone marrow cells (500,000) from symptomatic *Idh2^Lox-R140Q^ Flt3^GL-ITD^* mice were transplanted into lethally irradiated recipients with 200,000 Cd45.1 WT bone marrow. Line chart depicts WBC (K/μL; left) and hematocrit (%; right) at the indicated time point post transplantation. (n=17), Error bars reflect the mean ± s.e.m.; and p-values are calculated by Fisher’s LSD test (e), *\*\* p ≤ 0.01 *** p ≤ 0.001*.

**Extended Data Fig. 8:**
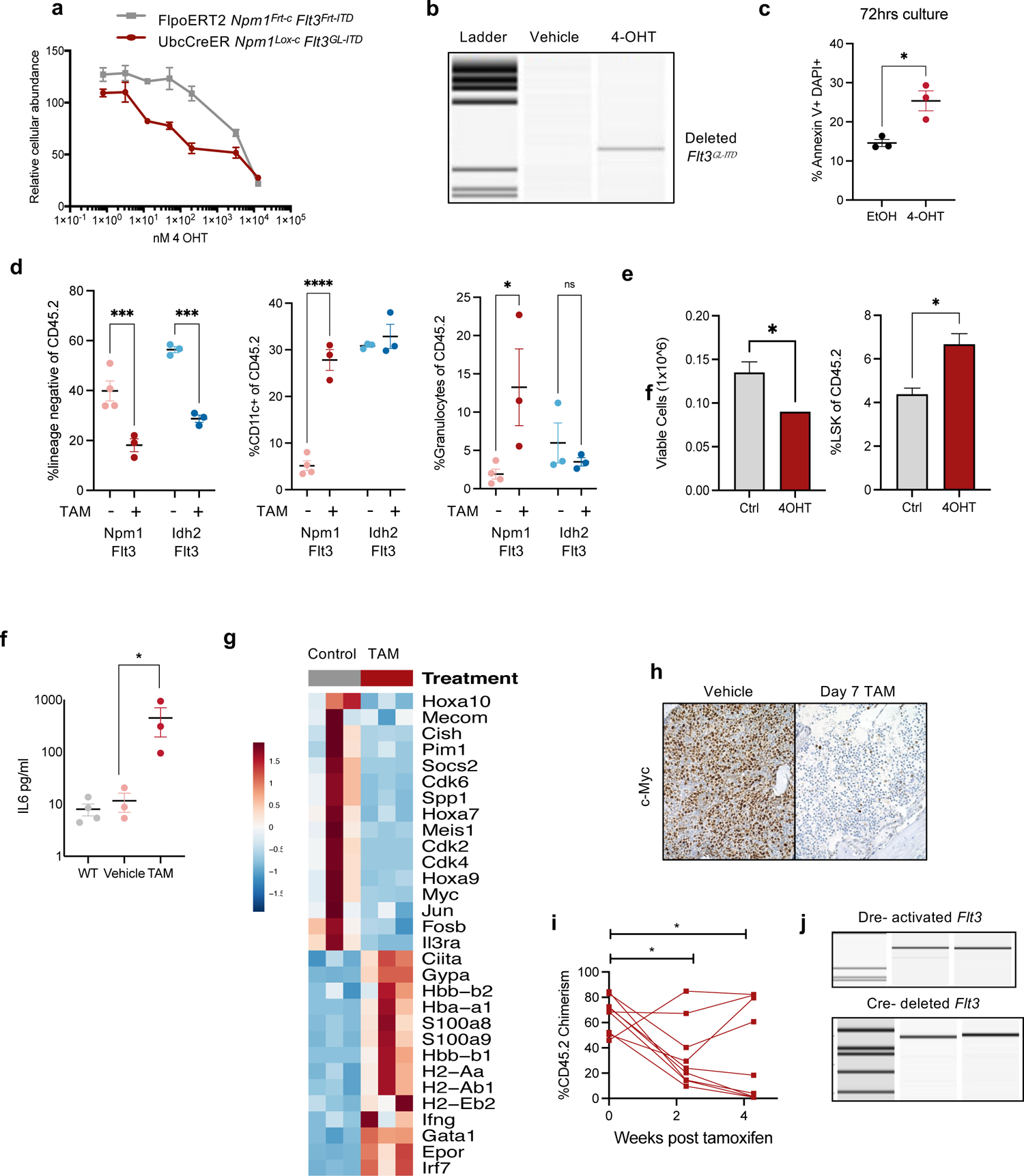
Immunophenotypic and molecular characteristics of *Flt3*-genetic ablation. **a,** cKIT+ cells from Rosa26:FlpoERT2 *Npm1^Frt-C^ Flt3^Frt-ITD^* (grey) and Ubc:CreER Vav1:DreStaPL-ODC *Npm1^Lox-C^ Flt3^GL-ITD^* (red) were cultured in Stemspan supplemented with 20ng SCF and treated with escalating doses of 4-OHT. Relative cellular abundance was assessed with PrestoBlue at 72 hours of culture. (n=3 per group). **b,** PCR depicting *Flt3^GL-ITD^* deletion with 4-OHT treatment *ex vivo*, visualized by capillary electrophoresis. **c,** Cells were cultured as in (a) with either 400nM 4-OHT or vehicle control. Apoptosis was assessed by Annexin V and DAPI by flow cytometry. (n=3 per group). **d,** Dot chart depicting %lin^-^ (left), %Cd11c^+^ (middle) and granulocytes (Cd11b^+^Gr1^high^; right) of Cd45.2 cells in bone marrow from control and TAM treated mice for the indicated disease models. (n=3-4 per group). **e,** cKIT+ cells from leukemic *Npm1^Lox-C^ Flt3^GL-ITD^* mice were plated over a bone marrow endothelial feeder layer and treated with 4-OHT (400nM) or vehicle control. Total cell number (left) and %LSK of Cd45.2 (right) were assessed 6 days after plating. (n=3 per group). **f,** Serum cytokine levels for IL-6 from WT, untreated *Npm1^Lox-C^ Flt3^GL-ITD^,* and TAM-treated (7d) *Npm1^Lox-C^ Flt3^GL-ITD^* mice. (n=3-4 per group). **g,** Row-normalized heatmap of RNA-sequencing data in untreated (grey) or 7 day TAM-treated (red) leukemic *Npm1^Lox-C^ Flt3^GL-ITD^* mice. **h,** Immunohistochemical detection of c-MYC in bone marrow from control and TAM (7 day) treated *Npm1^Lox-C^ Flt3^GL-ITD^* (400X). **i,** Peripheral blood chimerism of (%Cd45.2) of *Npm1^Lox-C^ Flt3^GL-ITD^* following the initiation of TAM-treatment. (n=9 per group). **j,** PCR evaluating Dre-mediated inversion (top) and Cre-mediated deletion (bottom) of *Flt3^GL-ITD^* in recurrent *Npm1^Lox-C^ Flt3^GL-ITD^* leukemias following TAM treatment. PCR products were visualized by capillary gel electrophoresis (Qiaxcel). (n=2 representative of a cohort of 9). Error bars reflect the mean ± s.e.m.; and p-values are calculated by Student’s T-test (b,d) and Fisher’s LSD test (c,e,i,j), *\* p ≤ 0.05 ** p ≤ 0.01* **** *p ≤ 0.0001*.

## Methods

### Mouse model generation

All protocols were approved by the MSKCC IACUC under protocol 07-10-016. Ubc:CreERT2 (strain 007001), RC::RLTG (strain 026931) and RC::FLTG (strain 026932) mice were purchased from Jackson Labs. The Rosa26:FlpoERT2 and Npm1^Frt-c^ mice were described previously and provided by the Trowbridge laboratory^9^. The Dnmt3a^Lox-R878H^, Npm1^Lox-c^ and *Idh2^R140Q^* were described previously ^40,41,42^. The *Flt3^Frt-ITD^* and *Flt3^GL-ITD^* mice were generated at Ingenious targeting labs. The targeting construct for *Flt3^Frt-ITD^* was generated A 9.8 kb genomic DNA used to construct the targeting vector was first subcloned from a positively identified C57BL/6 BAC clone (RP23-280N16). The region was designed such that the long homology arm (LA) extends ∼7 kb 5’ to the 5’ FRT cassette, and the short homology arm (SA) extends about 2 kb 3’ to the insertion of the inversion cassette. The inversion cassette is flanked by mutant F3 sites that point away from each other. The 3’ FRT site is placed right before the 3’ F3 site. The inversion cassette consists of exon 13-mutant exon 14* (humanized Flt3-IDT)-exon 15 and the flanking genomic sequences from upstream of exon 13 to downstream of exon 15 for correct splicing (Inv.saE13-14*-E15Sd). This cassette was inserted in the reverse direction downstream of exon 15. The Lox2272-flanked Neo cassette was inserted immediately upstream of the inversion cassette and is 235 bp away from wt exon 15. The targeting region is 985 bp containing exons 13-15. Ten micrograms of the targeting vector was linearized and then transfected by electroporation of C57Bl/6 (B6) embryonic stem cells. After selection with G418 antibiotic, surviving clones were expanded for PCR analysis to identify recombinant ES clones. After successful clone identification, the neomycin cassette was removed with a transient pulse of Cre recombinase and clones were reconfirmed following expansion. Finally, ES cells were injected in C57B6 mice via tetraploid complementation (NYU).

The targeting construct for *Flt3^GL-ITD^*, was generated as follows: A 9.8 kb genomic DNA used to construct the targeting vector was first subcloned from a positively identified C57BL/6 BAC clone (RP23-280N16). The region was designed such that the long homology arm (LA) extends ∼5.7 kb 5’ to the cluster of Lox2272-Rox-Rox12-Lox2272 sites, and the short homology arm (SA) extends about 1.9 kb 3’ to the Neo cassette and 3’ Rox12 site. The inversion cassette is in between the second set of Lox2272 and Rox sites, and it consists of exons 12-15 with the IDT mutation engineered in exon 14 (Flt3-IDT) and the flanking genomic sequences from upstream of exon 12 to downstream of exon 15 for correct splicing (SaE12-W51-15Sd). The inversion cassette replaces wt exons 12-15 and the same flanking genomic sequences included in the cassette. The deleted region is 2.19 kb. The FRT-flanked Neo cassette is inserted immediately downstream of the inversion cassette. Each pair of the recombination sites are in opposite direction. Ten micrograms of the targeting vector was linearized and then transfected by electroporation of FLP C57Bl/6 (BF1) embryonic stem cells. After selection with G418 antibiotic, surviving clones were expanded for PCR analysis to identify recombinant ES clones. The Neo cassette in targeting vector has been removed during ES clone expansion. Targeted iTL BF1 (C57BL/6 FLP) embryonic stem cells were microinjected into Balb/c blastocysts. Resulting chimeras with a high percentage black coat color were mated to C57BL/6 WT mice to generate Germline Neo Deleted mice. Tail DNA was analyzed as described below from pups with black coat color.

The knockin constructs for the *Vav1:DreStaPL*-ODC and H11:CAG:DreStaPL-ODC mice were synthesize (IDT) and assembled via Gibson assembly. Knockin constructs were injcted into zygotes with Cas9 ribonucleotide complexes targeting either Vav1 or H11 at Memorial Sloan Kettering Cancer Center using the Mouse Genetics Core Facility. Founders were screened by PCR genotyping. Complete sequences of all mouse models will be available on benchling.

#### Tissue harvest and bleeds

Peripheral blood was isolated by submandibular bleeds. Complete blood counts were assessed on a ProCyte by IDEXX. For flow cytometry analysis whole blood was lysed with RBC lysis buffer (Biolegend). For terminal tissue isolation, mice were euthanized with CO2 asphyxiation, tissues were dissected and fixed with 10% Zinc formalin for histopathological analysis. For whole bone marrow was isolation, the femur, hip and tibia were dissected and cleaned. Cells were isolated by centrifugation at 8000xG for 1 minute. Single cell suspension were generated from crushed whole spleen and filtered through a 70uM filter. RBC lysis was performed (Biolegend) and cells were prepared for downstream processing or frozen in 10%DMSO+90% fetal bovine serum.

#### Bone marrow transplant

Donor cells were either prepared fresh or thawed from frozen cells. Fresh cells were prepared as described above. When prepared from frozen cells, cells were thawed at 37C into warm FACS buffer (PBS+2%fetal bovine serum (FBS)), centrifuged at 1500rpm for 5 minutes, washed once with warm and resuspended in PBS. Donor cells and support cells were mixed at desired ratios as indicated. The day before injection, Cd45.2 recipient mice were irradiated with 900cGy using an Xcellerator cesium irradiator. Recipient mice were heated under a heat lamp for 5-10 minutes to induce vasodilation in preparation for tail vein injection. Mice were restrained using a rotating tail injector restrainer and 200ul of cells were injected per mouse.

#### In vivo drug dosing

Tamoxifen (TAM) powder (500mg) was dissolved in 25ml of corn oil, wrapped in foil and shaken at 225rpm overnight. TAM was the aliquoted and stored at −80C. Upon dosing, TAM was thawed and dosed at 200ul (4mg) per mouse via oral gavage. Tamoxifen chow was purchased from Envigo and provided *ad libitum*. Grazoprevir (GZV) powder (100mg) was dissolved in 1ml of DMSO and frozen at −80C. GZV was administered *in* vivo with Ritonavir (RTV) and Lopinavir (LPV). RTV 40mg/ml DMSO) and LPV (40mg/ml DMSO) were mixed together in a 1:4 ratio (RTV:LPV) and stored at −80C. For *in vivo* dosing, 27.5μL GZV was added to 27.5 RTV/LPV, 400μL of PEG400, and finally 635μL PBS. Mice were dosed with the GZV+RTV+LPV mix at 200ul via intraperitoneal (IP) injection with a 27.5G insulin syringe. Trimethoprim (TMP) powder (100mg) was dissolved in 1ml of DMSO and frozen in 110ul aliquots at −80C. Upon dosing, TMP was made fresh by thawing an aliquot and adding 440ul of PEG400 and 550ul of PBS to get a final volume of 1100ul. Mice were dosed at 200ul via intraperitoneal (IP) injection with a 27.5G insulin syringe.

#### Cloning

Codon optimized versions of Cre, CreERT2 and Dre were synthesized (Thermo) and subcloned into pENTR using the directional TOPO cloning kit. Flpo was PCR amplified from a FlpoERT2 construct provided by Dr. Alex Joyner at MSKCC, and then TOPO cloned into pENTR using the same approach. DreStaPL and DreStaPL-ODC were generated by a gibson reaction with the backbone generated through inverse PCR of the pENTR vector, and the StaPL containing insert synthesized as a gBlock for IDT. A similar approach was used for inserting DHFR downstream of Flpo. These pENTR constructs were then cloned into a MSCV:IRES GFP destination vector using a gateway cloning approach (Thermo: LR Clonase II).

#### Virus generation and viral infection

MSCV retroviral vectors were transfected with pCL-Eco packaging vector into 293T-17 cells using JetPRIME following the manufacturers protocol (5ug each, 1:1). Conditioned media was collected 72 hours later and passed through a 0.45uM filter. Virus was either used fresh or stored at −80C for later usage. For viral infection in Figure 4 and Extended Data 8, lineage negative bone marrow was isolated using the Mouse Hematopoietic Progenitor Cell Isolation Kit (StemCell 19856) and plated over retronectin coated 6 or 12 well plates for 2 hours in RPMI+10%FBS supplemented with Penicillin/Streptomycin, 20ng SCF, 10ng IL6 and 10ng IL3. Virus and 100X HEPES was added to the wells, then plates were spun for 1 hour at 800xG at 37C. Following spin, small molecule ligands used to activate recombinases were added to a final concentration as indicated ethanol (v/v 0.0015%), DMSO (v/v 0.1%), 4-hydroxytamoxifen (4-OHT; 400nM), trimethoprim (TMP; 1uM) or grazoprevir (GZV, 10uM). Cells were harvested for flow cytometric analysis 48 hours post infection. For viral infection followed by transplantation in Extended Data 8, a similar approach was utilized with the following modifications. Donor mice were treated with 5-FU 6 days prior to harvest. Mice were euthanized as described above, and bone marrow was isolated by flushing bones with 10ml of FACS buffer with a 26.5 gauge needle. Whole bone marrow was plated retronectin plates as described above, and cells were harvested for transplant 2 hours after infection.

#### Histology staining and immunohistochemistry, photography

Spleen, liver and tibia samples were fixed (4% paraformaldehyde) for >24 hours and embedded in paraffin. Sections were cut using microtome (Mikrom International AG), mounted on slides (ThermoScientific), and dried at 37°C overnight. Hematoxylin and eosin (H&E) staining was performed on a COT20 stainer (Medite). All sample handling and preparation was performed by Tri-Institutional Laboratory of Comparative Pathology (LCP) facility. The following antibodies were used for immunohistochemistry: phospho-44/42 MAPK (Erk1/2) (Cell Signaling 4376, 1:100). Pictures were taken at 400X magnification using an Olympus microscope and analyzed with Olympus Cellsens software. Tissue sections were formally evaluated by a hematopathologist (W. Xiao).

#### Cell culture

For Extended Data 8a, 5000 cKIT+ cells from either Ubc:CreER *Npm1^Lox-c^Flt3^GL-ITD^* or *Rosa26:FlpoERT2 Npm1^Frt-c^Flt3^Frt-ITD^* leukemias were added to non TC coated 96 well plate in StemSpan (STEMCell) supplemented with Penicillin/Streptomycin and 20ng/ml of SCF with a titration range of 4-OHT. Cellular abundance was read out 72 hours later using the Presto Blue assay (Thermo) on a Cytation 3 (BioTek) plate reader. Similar cultures conditions were used in Extended Data 8b for apoptosis analysis, scaled to a 12 well plate with 400nM 4-OHT. Cell viability was assessed with tryphan blue on a Vicell Blue Cell Counter (Beckman Coulter).

*Ex vivo* cultures over bone marrow endothelial cells were performed as previously described. Briefly, 100,000 myristoylated-Akt immortalized bone marrow endothelial cells were plated into a fibronectin coated 12 well plate for 48 hours. Cultures were washed twice with PBS, and then 50,000 cKIT+ splenocytes from Ubc:CreER *Npm1^Lox-c^Flt3^GL-ITD^* leukemic mice were plated in StemSpan with Penicillin/Streptomycin and 20ng/ml of SCF. Media was changed 3 days later, and on day 6, cultures were harvested with Accutase (Biolegend) for viability by tryphan blue and assessment by flow cytometry.

#### Flow cytometry and sorting

Freshly isolated or cryopreserved cells were washed twice with FACS buffer (phosphate buffered saline (PBS)+ 2% fetal bovine serum). Cells were incubated with antibody cocktails for 15 minutes at 4°C. Complete antibody details can be found in **Supplementary Table 2**. Following antibody incubation, cells were washed with FACS buffer and resuspended in a DAPI containing FACS buffer solution for analysis and sorting. Flow cytometry analysis was performed on a BD LSRFortessa with analysis using FACSDiva and FlowJo (v10.9). Cell isolation with FACS was performed on a Sony SH800.

#### Nucleic acid isolation and RNA-sequencing library generations

DNA was isolated using the DNeasy Blood & Tissue Kit (Qiagen). RNA was isolated using the Direct-zol RNA Microprep Kit (Zymo Research, R2061) and quantified using the Agilent High Sensitivity RNA ScreenTape (Agilent 5067-5579) on an Agilent 2200 TapeStation. 3’ multiplexed RNA-sequencing was performed with the Takara SMART-Seq v4 3’ DE Kit (Takara 635040) followed by Nextera XT (Illumina FC-131-1024) library preparation. For full length RNA-sequencing, cDNA was generated from 1ng of input RNA using the SMART-Seq HT Kit (Takara 634455) at half reaction volume. cDNA and tagmented libraries were quantified using High Sensitivity D5000 ScreenTape (5067-5592) and High Sensitivity D1000 ScreenTape respectively (5067-5584). Libraries were sequenced on a NovaSeq at the Integrated Genomics Operation (IGO) at MSKCC.

#### Mass Cytometry and data analysis

Cryopreserved mouse whole bone marrow or spleen cells were thawed and counted. Approximately 1 million from each mouse analyzed were used. Live cells were rested for 2 hours in RPMI + 10% FBS (Gibco) (cRPMI) and then stained for viability with 5 M Cell-ID Cisplatin (Fluidigm 201198, 201194) for 2.5 minutes and washed in cRPMI. Cells were fixed with paraformaldehyde at a final concentration of 1.6% for 10 mins at RT in the dark and washed with Maxpar PBS (Fluidigm). Palladium mass-tag barcoding was performed as previously described using combinations of 6 palladium isotopes with Cell-ID 20-plex Pd barcoding kit (Fluidigm).1,2 All barcoded cells were then combined and washed with Maxpar cell staining buffer (CSB) (Fluidigm) and stained first with anti-CD16/32-159Tb for 30min, followed by the rest of the surface antibody cocktail for 30 mins (**Supplementary Table 2**). Cells were then washed with PBS, and permeabilized with ice cold 100% methanol (Fischer Scientific) for at least 30 mins. Following permeabilization, cell were washed and pelleted twice with CSB followed by intracellular antibody staining for 30 mins. Just prior to data collection, cells were stained with 250 nM Cell-ID Iridium intercalator (Fluidigm) in PBS with 1.6% PFA for 30 mins at 4°C and then washed per protocol. Cells were then washed and rehydrated in double deionized water and collected on a Helios mass cytometer (Fluidigm). Barcorded .fcs files were normalized and debarcoded with Fluidigm Debarcoder software. Files were downloaded and gated in Cytobank (BeckmanCoulter) for single intercalator positive cells. Cd45.2-positive cells were gated and files were downloaded and loaded into R for use by the CATALYST package.3,4 Analysis and plots were created using this following standard workflows.

#### mRNA electroporation

Lineage negative bone marrow cells were isolated using the Mouse Hematopoietic Progenitor Cell Isolation Kit (StemCell 19856) and electroporated with a custom Dre mRNA (Tri-Link) using the Thermo Neon electroporation device according to the manufacturer’s protocol (100ul kit, Thermo MPK10025). In brief, lineage negative bone marrow cells were resuspended in 153ul of buffer T and 17ul of Dre mRNA (1ug/ul) was added and mixed by pipetting. Immediately after RNA addition, cells were electroporated using one pulse of 1700V for 20ms. Following electroporation, cells were added to serum free media (StemSpan) with 50ng SCF and 10ng IL3 without antibiotics. For transplantation experiments, cells were incubated at 37 degrees for 1 hour and then injected into mice as described above. For extended culture, antibiotics were added 4 hours after electroporation.

#### Bioinformatic analysis

*RNA-sequencing:* FASTQ files were demultiplexed using a java script from Takara. FASTQ files were mapped and transcript counts were enumerated using STAR (genome version mm10 and transcript version genecode M13). Counts were input into R and RNA-sequencing analysis was completed using DESeq2. Gene set enrichment analysis was performed using the fgsea package with genesets extracted from the msigdbr package. Single sample gene set enrichment analysis was performed using the gsva package. Figures were generated using ggplot2 and tidyheatmaps packages. Complete scripts will be made available on github.

#### Patient samples

Patients with acute myeloid leukemia between 2014-2020 were studied. Informed consent from patients was obtained in accordance with the Declaration of Helinski and according to protocols by the institutional IRBs. This study was approved by MSKCC IRB (protocol #15-017) and Thomas Jefferson University (TJU) IRB (protocol #17D.083). Diagnosis, AML status, and normal karyotype were confirmed and assigned based on World Health Organization classification criteria. Samples from patients were collected and process by the Human Oncology Tissues Bank at MSKCC or the Heme Malignancy Repository at TJU. Bone marrow mononuclear cells were isolated by Ficoll and viably frozen. MSKCC samples were subjected to high-throughput genetic sequencing with HemePACT, a targeted deep sequencing assay of 685 gene recurrently mutated in hematologic malignancies. Variants and short insertion/deletions are identified as described previously (Miles, Bowman Nature 2020). Samples were selected from patients harboring a confirmed FLT3-ITD mutation co-mutated with DNMT3A and/or NPM1 mutations where 1) all mutations were covered by the Mission Bio Custom amplicon panel, 2) variant allele frequency of each mutation was >5%, and 3) cell number collected was feasible for downstream processing (>5×10^6^ cells).

#### Single-cell DNA+Protein sequencing library preparation and sequencing

Patient samples were thawed and quantified using a Countess cell counter. Viable cells (1.0-4.0×10^6^) were resuspended in Cell Staining Buffer (Mission Bio) and incubated with TruStain FcX and Blocking Buffer (Mission Bio) for 15min on ice. The Biolegend TotalSeq-D Heme Panel consisting of 45 oligo-conjugated antibodies (AOC; Supplementary Table 2) and a custom TotalSeq-D AOC against human CD135 (0.5mg/mL; FLT3/FLK2; clone BV10A4H2) was resuspended in Cell Staining Buffer and then incubated with the blocked cell suspending for 30 min on ice. Cells were washed multiple times with Cell Staining Buffer and finally resuspended in Cell Buffer (Mission Bio). Stained cells were requantified, loaded into a Tapestri microfluidics cartridge, encapsulated, lysed and barcoded as previously described (Miles, Bowman Nature 2020, Pellegrino, M. Genome Res, 2018). DNA and Protein PCR products were then isolated and purified as previously described (Miles Bowman Nature 2020). All libraries were quantified using an Aligent Bioanalyzer, normalized, and pooled for sequencing on an Illumina NovaSeq by the MSKCC Integrated Genomics Core. FASTQ files were processed via the Tapestri pipeline on the Mission Bio cloud server. H5 files were downloaded and processed with in house scripts in R, complete scripts and tutorial wil be made available on github. Briefly, H5 files were read into using the h5read function from the ‘rhdf5’ package. Variants were filtered if they were not genotyped in less than 20% of cells, or possessed an initial variant allele frequency <0 .005%. Subsequently, non-synonymous protein encoding variants were filtered with a depth (DP) cutoff of 10, gene quality (GQ) cutoff of 30, and allele frequency cutoff of <20% for wildtype, 20-80% for heterozygous and >80% for homozygous calls. Cells that possessed GATK calls from the tapestri pipeline that passed these filtered were retained, and were consolidated into isogenic clones. Protein data was generated from the tapestri pipeline as raw counts, and imported into R as a Seurat object. The protein data was logNormalized, centered and scaled on a per sample basis, and the top 4 principle components were used to as input in the KNN neighbor identification. Subsequent community identification was performed with the ‘FindClusters’ function with a resolution variable of 0.25. The clone information was applied as metadata, and all subsequent analysis occurred in Seurat, including generation of ridgeplots, feature heatmaps, and UMAPs.

## Data availability

All raw and processed sequencing data will be made accessible via the NCBI Gene-Expression Omnibus (GEO) mouse and dbGAP.

## ACKNOWLEDGEMENTS

We are grateful to members of the Levine Lab for their discussion of the work. We would also like to acknowledge Dr. Alex Joyner (MSKCC), Dr. Patricia Jensen (NIH) and Dr. Tudor Badea (NIH) for discussion on Dre-Rox technology and Dr. George Vassiliou for sharing the *Npm1^Lox-C^* mice. This work was supported by National Cancer Institute awards P01 CA108671 (R.L.L.). R.L.L. was supported by a Leukemia and Lymphoma Society Scholar award. R.L.B. was supported by a Damon Runyon-Sohn Fellowship and the National Cancer Institute (K99CA248460). A.J.D. is a William Raveis Charitable Fund Physician-Scientist of the Damon Runyon Cancer Research Foundation (PST-24-19). He also has received funding from the National Institute of Health (T32CA009207), American Association of Cancer Research (17-40-11-DUNB) and the American Association of Clinical Oncology. L.A.M. was supported by the National Cancer Institute (K99CA252005). Studies supported by MSK core facilities were supported in part by MSKCC Support Grant/Core Grant P30 CA008748 and the Marie-Josée and Henry R. Kravis Center for Molecular Oncology. R.L.L. is also supported by a Leukemia & Lymphoma Society Specialized Center of Research grant.

## AUTHOR CONTRIBUTIONS

R.L.B., A.D, and R.L.L. conceived and designed the study. R.L.B., A.D., T.M., M.R.W., I.F.M., S.F.C., P.S.V., and P.B.F., designed and executed experiments. T.M., S.E.E., S.M., L.C.,Y.P., A.M.B, I.S.C., D.L., N.B., and A.K. provided technical support on experiments. R.L.B., A.J.D. W.X., M.R.W., I.F.M., S.C., and P.B.F. analyzed the data W.X., provided histopathological assessment. R.L.B. performed all computational analysis. C.R.P., M.T.J., and P.B.J. designed, executed, and analyzed the CyTOF experiments. S.M., M.P.C., L.A.M., P.B.F., and J.J.T. provided critical discussion on experimental design and crucial reagents. R.L.L. supervised the study. R.L.B. and R.L.L. wrote the manuscript with significant revisions and critical feedback from A.D. and L.A.M., all authors reviewed and commented on the final manuscript.

